# Copy Number Variation: A Substrate for Plant Adaptation and Stress Response in Arabidopsis

**DOI:** 10.1101/2025.08.15.670553

**Authors:** Piyal Karunarathne, Yannick Schäfer, Anna Glushkevich, Alison D. Scott, Leen Abraham, Juliette de Meaux, Polina Yu Novikova, Thomas Wiehe, Laura E. Rose

## Abstract

Copy number variation (CNV) is a major source of genomic diversity that shapes gene family evolution and may contribute to ecological differentiation, yet its genome-wide ecological relevance remains poorly understood. Here, we analyzed CNV across four Brassicaceae species (*Arabidopsis thaliana, A. lyrata, A. halleri*, and *Arabis alpina*) to identify gene families undergoing rapid expansion or contraction. Using a birth–death model, we identified 231 rapidly evolving gene families spanning diverse functional categories. We then characterized CNV within populations of *A. thaliana* and *A. lyrata* using population-scale long-read assemblies. CNV exhibited strong heterogeneity across gene families and species, with contrasting evolutionary outcomes: *A*. thaliana showed greater retention of duplicated copies, whereas *A. lyrata* exhibited higher pseudogenization and turnover. CNV profiles were strongly structured geographically, reflecting known demographic and phylogeographic patterns in both species. Environmental association analyses revealed species-specific architectures: CNV in *A. thaliana* showed a diffuse, polygenic association with climate, whereas in *A. lyrata* associations were more strongly coupled to population structure. Despite the functional diversity of rapidly evolving families, environmentally associated CNVs were significantly enriched in defense- and stress-related functions. These results demonstrate that CNV–environment relationships emerge at the level of gene family networks and are shaped by genomic architecture and lineage history, highlighting CNV as a context-dependent driver of genome evolution and ecological differentiation.

**Summary:** Gene copy number variation—where individuals carry different numbers of copies of the same gene—is a potentially important but poorly understood source of genetic diversity. The authors studied how copy numbers evolve across four mustard family plant species and whether this variation helps plants adapt to different environments. They found that gene families involved in disease resistance and stress responses were especially prone to rapid copy number changes. These changes tracked geography and local climate, but in ways that differed between species depending on their evolutionary history. The findings highlight gene copy number variation as a key, underappreciated contributor to how plants adapt to their environments.

## 1 Introduction

Gene duplication and gene loss are fundamental evolutionary processes that generate genetic diversity across genomes and lineages. Among the different forms of structural genomic variation, copy number variation (CNV) represents a major source of heritable genomic variation, defined as insertions, deletions, or duplications of genomic segments ranging from kilobases to megabases in size (Freeman et al., 2006; Redon et al., 2006). CNVs can encompass entire genes or gene clusters, and influence phenotypic variation through altered gene dosage, expression levels, or regulatory interactions (Zhang et al., 2009; Schrider and Hahn, 2010). In plants, CNVs have been implicated in a wide range of adaptive traits, including disease resistance, flowering time regulation, and responses to abiotic stress (Maron et al., 2013; Cao et al., 2011; Prunier et al., 2017).

The importance of CNVs in plant adaptation is increasingly recognized, particularly in relation to biotic and abiotic stress responses. Several gene families involved in pathogen recognition (e.g. NBS-LRR genes), detoxification, or signaling have been shown to exhibit dynamic CNV patterns both within and between species (Kang et al., 2012; Rizzon et al., 2006; Hanikenne et al., 2008). Among species, these gene families can often be described by a birth-and-death model of evolution, whereby gene duplication is followed by functional diversification, neofunctionalization, or pseudogenization, while other paralogs are lost (e.g., Nei and Rooney, 2005; also see Otto and Wiehe, 2023 for an alternative mechanism). However, at the intraspecific level, duplicated genes can also retain equivalent biochemical functions, with variation in copy number modulating gene dosage and adaptive responses (e.g., *HMA4* genes in *Arabidopsis halleri* –Hanikenne et al., 2008). While the population-level consequences of CNV have been explored in domesticated crops (Muñoz-Amatriaín et al., 2013; Ma et al., 2025), fewer studies have examined the evolutionary and ecological role of CNV across multiple closely related species in natural populations.

The Brassicaceae family, and particularly the *Arabidopsis* genus, provides an ideal system to investigate the evolution and ecological consequences of copy number variation (CNV). Highquality reference genomes and extensive population genomic resources are available for species such as *A. thaliana* and *A. lyrata*, enabling fine-scale analyses of structural variation and its environmental associations (Alonso-Blanco et al., 2016; Hu et al., 2011; Novikova et al., 2016). These species span broad geographic and ecological ranges and differ markedly in life history and demographic history, offering a powerful comparative framework. *A. thaliana*, a predominantly selfing annual, exhibits strong population structure shaped by glacial history, whereas *A. lyrata*, an outcrossing perennial with a circumboreal distribution, represents a contrasting evolutionary context (Alonso-Blanco et al., 2016; François et al., 2008; Hämälä et al., 2018a). Together with the availability of long-read assemblies, population-scale genomic data, and environmental metadata, this system enables high-resolution investigation of how CNV contributes to gene family evolution and ecological differentiation. Previous studies have linked CNVs in specific gene families, such as NB-LRR and F-box genes, to adaptive responses in *A. thaliana* (Todesco et al., 2010; Exposito-Alonso et al., 2019), providing a foundation for extending these insights to a genome-wide, comparative context.

Here, we test the hypothesis that copy number variation (CNV) in gene families is non-randomly distributed across functional categories and shaped by both environmental gradients and lineage-specific genomic context, with distinct evolutionary outcomes following gene duplication. Specifically, we hypothesize that (i) rapidly evolving gene families are enriched for defense- and stress-related functions, reflecting selective pressures imposed by heterogeneous environments; (ii) CNV within these families shows structured associations with environmental variables after accounting for population structure; (iii) closely related species differ in the architecture of CNV–environment relationships due to differences in genome organization and demographic history; and (iv) duplicated gene copies follow divergent evolutionary fates, with lineage-specific balances between retention, functional divergence, and pseudogenization. To address these hypotheses, we combine comparative genomic analyses across Brassicaceae species with population-scale long-read data from *A. thaliana* and *A. lyrata* to identify rapidly evolving gene families, characterize their functional composition and genomic organization, assess the structural and functional fate of duplicated copies, and evaluate geographic and environmental patterns of CNV. This integrative framework provides a genome-wide perspective on how gene duplication and CNV contribute to gene family evolution and ecological differentiation in natural populations.

## 2 Materials and Methods

### 2.1 Analysis of gene family expansion and contraction

Gene families were inferred as orthogroups (the complete set of genes descended from a single gene in the last common ancestor) using OrthoFinder2 (Emms and Kelly, 2019), based on curated reference proteomes of four Brassicaceae species (*A. thaliana, A. lyrata, A. halleri, Arabis alpina*) with *Carica papaya* as the outgroup. Proteomes for *A. thaliana, A. lyrata, A. halleri*, and *C. papaya* were downloaded from the Phytozome database (Goodstein et al., 2012), and the *A. alpina* (V4) from *A. alpina* genome portal (http://www.arabis-alpina.org) (assembly details in Supplementary Table S0). OrthoFinder2 identified orthogroups and reconstructed gene and species trees, which were then used to define gene families for comparative analyses.

Rates of gene family expansion and contraction (i.e., gene family evolutionary rates [*λ*]) were estimated with CAFE 5 (Hahn et al., 2005; Mendes et al., 2020), which applies a birth–death model of gene gain and loss across a time-calibrated species tree. We tested multiple models of evolutionary rate variation (i.e., gamma [*γ*] model) and incorporated an error model (*ϵ*) to account for annotation artifacts. Gene families showing significant expansion or contraction (p < 0.01: “Rapidly-Evolving” families) were retained for downstream functional characterization and CNV analysis. While inference of rapidly evolving families can be affected by annotation quality and gene fragmentation, particularly in TE-rich regions, the use of curated reference proteomes and likelihood-based modeling in CAFE 5 helps distinguish true signals from artefacts; enrichment of functionally coherent categories further supports their biological relevance. Full methodological details are provided in Supplementary Section **S1**.

### 2.2 Gene family function assessment and categorization

We used *A. thaliana* genes to find the predicted functions of the members of all “rapidly-evolving” gene families. This was done using custom python and R scripts to search the gene function databases of UniProt (Consortium, 2022) and TAIR (Berardini et al., 2019). Based on the primary gene function, we categorized the genes into ten broader functional categories. We further categorized these gene families into two groups based on the functional similarity of their members. Gene families in which all members share the same annotated function were categorized as functionally “conserved” and others as functionally “diversifying.”

### 2.3 Assessment of CNV across rapidly evolving gene families

#### 2.3.1 Long-read *de novo* assembly compilation

To assess copy number variation (CNV) within species across rapidly-evolving gene families, we compiled a curated dataset of long-read *de novo* genome assemblies comprising 137 *A. thaliana* assemblies from NCBI (Tab. S3) and 24 *A. lyrata* assemblies (Tab. S4) generated in this study and a complementary pangenome project. The *A. thaliana* dataset integrates assemblies produced using both PacBio CCS and Oxford Nanopore (ONT) technologies, whereas the *A. lyrata* dataset consists exclusively of PacBio CCS assemblies (see Supplementary Information S2 for assembly details).

Assembly quality was evaluated using BUSCO and standard assembly metrics. To assess potential technical biases, we examined the effects of sequencing technology and assembly quality on CNV estimates. Three quality metrics—BUSCO completeness, contig N50, and total assembly size—were retained as technical covariates in downstream analyses of CNV and CNV–environment associations (see Tab. S3&4).

#### 2.3.2 CNV detection

Gene copies of all rapidly-evolving gene families were identified using a three-step BLAST-based procedure optimized for long-read *de novo* assemblies. First, reference gene sequences from each gene family were queried against each assembly using `blastn` (Camacho et al., 2009), allowing local alignments to capture full-length as well as partial matches. Second, BLAST hits with ≥ 90% sequence identity were retained and overlapping hits were merged based on their genomic coordinates, provided that the merged region did not exceed 1.3× the reference gene length. Third, in regions where multiple closely related genes produced overlapping hits, only the gene with the highest sequence identity was retained. To exclude short fragments and spurious matches (e.g., arising from transposable elements), only regions covering ≥50% of the reference gene length were considered.

This approach leverages long-read *de novo* assemblies, which provide high-resolution representation of structural variation and gene copy architecture, including complex rearrangements and tandem duplications that are difficult to resolve with short-read data. Within this framework, the BLAST-based strategy offers a straightforward way to recover full-length, partial, and structurally divergent copies at the locus level. It ensures that all detectable copy variants are captured within the genomic context of each gene family, making it well suited for targeted, gene family–focused CNV inference (detailed pipeline and limitations are in Suppl. Info. S3).

#### 2.3.3 Data restructuring and summary statistics

Copy number variation was assessed across all rapidly evolving gene families in *A. thaliana* and *A. lyrata* using species-specific gene family × assembly copy-number matrices generated from the assembly-based CNV detection pipeline described above. In these matrices, rows correspond to gene families and columns to assemblies, with each cell containing the total copy number observed for a given gene family in a given assembly.

For each gene family within each species, we calculated descriptive statistics including mean copy number, variance, and coefficient of variation (CV). These statistics were used to summarize the extent of within-species copy-number heterogeneity across assemblies and to identify families with non-zero copy-number variance. Additional family-level summaries, including minimum, maximum, median, standard deviation, and dispersion (variance/mean), were calculated for downstream comparative visualization.

#### 2.3.4 Within-species and between-species comparison of copy-number variation

To characterize broad patterns of CNV across the full dataset, we fitted generalized linear mixed models using the `glmmTMB` package (McGillycuddy et al., 2025). Because copy number is a count variable and showed overdispersion, models were fitted with a negative binomial error distribution. First, a null model was fitted to estimate background variation in copy number while accounting for repeated observations across both gene families and assemblies: *copy* ~ *1 + (1* | *family_id) + (1* | *assembly)*. This model captures overall copy-number heterogeneity while accommodating non-independence among observations belonging to the same gene family or assembly. Second, to test whether copy-number patterns differed between species, we fitted a model including species as a fixed effect: *copy* ~ *species + (1* | *family id) + (1* | *assembly)*. The two models were compared using likelihood-ratio tests. A significant improvement of the species model over the null model was interpreted as evidence that mean gene family copy number differs between *A. thaliana* and *A. lyrata* at the global level.

To assess species differences at the individual gene-family level, we performed family-wise Wilcoxon rank-sum tests comparing copy-number distributions between species. For each family, we also calculated the mean copy number in each species and the log_2_ fold change:

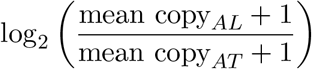

where 1 was added to both means to avoid undefined values for families with zero copies in one species. Resulting *P*-values were corrected for multiple testing using the Benjamini–Hochberg false discovery rate (FDR) procedure.

#### 2.3.5 Ordination of multivariate CNV profiles

To visualize major axes of variation in gene family copy-number profiles among assemblies, we performed principal component analysis (PCA) on the family × assembly copy-number matrices under three complementary transformations.

First, we performed a *raw PCA* on log-transformed copy numbers using `log1p` transformation, with variables scaled prior to ordination. This analysis retains the overall magnitude of among-family differences in copy number while reducing the influence of highly duplicated families. Second, we performed a *within-species standardized PCA*. For this analysis, log-transformed copy-number matrices were standardized within each species before being combined, thereby emphasizing relative CNV profiles within species while reducing the contribution of species-wide differences in total copy number. Third, we performed a *family-centered PCA* by centering log-transformed copy numbers by gene family across all assemblies without variance scaling. This transformation emphasizes deviations from family-specific average copy number and facilitates visualization of relative expansion or contraction patterns among assemblies. These three PCA approaches were applied to the combined dataset and, separately, to each species-specific matrix. Species-specific PCAs were further interpreted in relation to available population metadata.

#### 2.3.6 Multivariate testing of species differentiation

To test whether multivariate CNV profiles differed between species, we calculated Euclidean distances from the within-species standardized combined matrix and applied permutational multivariate analysis of variance (PERMANOVA) using `adonis2` from the `vegan` package (Oksanen et al., 2022), with species as the grouping factor. Because significant PERMANOVA results can arise either from differences in centroid position or from unequal within-group dispersion, we additionally tested homogeneity of multivariate dispersion using `betadisper`. Together, these analyses allowed evaluation of whether *A. thaliana* and *A. lyrata* differ in overall CNV structure across the 231 gene families.

### 2.4 Transposable element (TE) landscape around gene families

To assess the association between transposable elements (TEs) and gene family copy number variation, we quantified TE content around all gene family loci across all long-read assemblies of *Arabidopsis thaliana* and *A. lyrata*, and compared it to single-copy genes. TE annotation was performed using `RepeatMasker` (v4.1.5; Smit et al., 2013) with species-specific repeat libraries compiled from Repbase (Bao et al., 2015), Dfam (Hubley et al., 2016), and curated *Arabidopsis* resources (Hu et al., 2011).

Gene coordinates for gene family members were obtained from the CNV analysis described above. As a reference set, we defined single-copy genes by selecting genes with copy number < 2 across assemblies. Analyses focused on major TE classes known to be associated with CNV (Morgante et al., 2005; Kapitonov and Jurka, 2007), including DNA transposons (e.g., Helitron, MULE-MuDR), long terminal repeat (LTR) retrotransposons (Copia, Gypsy, Ty3), and non-LTR elements (LINEs and SINEs).

To characterize TE distribution around genes, we quantified TE content across a series of nested genomic windows centered on each gene: gene body (0 bp), ±1 kb, ±10 kb, ±25 kb, and ±50 kb. For each gene and window, we calculated: (i) number of TE insertions (*n*_TE_), (ii) total TE-covered base pairs, (iii) TE fraction (proportion of the window occupied by TEs), and (iv) distance to the nearest TE. The significance was assessed separately for each window using Wilcoxon rank-sum tests. Specifically, we tested for differences in: (i) TE counts (*n*_TE_), (ii) TE fraction, and (iii) distance to nearest TE between gene family and single-copy genes.

These metrics were computed separately for each assembly and gene. This window-based approach allows evaluation of both local TE insertions within gene bodies and broader TE landscape structure in flanking regions. Together, these analyses provide a multi-scale characterization of TE association with gene family expansion and duplication across both species.

### 2.5 Structural variation and evolutionary fate of gene family members

We applied a three-step framework to assess the structural integrity and putative functional relevance of gene family members across assemblies.

**Step 1: Locus delineation and re-annotation**. For each reference gene, candidate homologous loci were delineated using the genomic interval spanning the minimum and maximum BLAST hit coordinates (as described in CNV detection above). Sequences were extracted with flanking regions (+25% of gene length on both sides) to capture complete coding regions. Each locus was subsequently re-annotated by aligning the reference protein sequence to the genomic region using `exonerate` (Slater and Birney, 2005) (protein2genome model), and coding sequences were reconstructed from predicted CDS coordinates.

**Step 2: Structural classification**. Structural integrity of each copy was evaluated based on coding sequence completeness and translation. Predicted proteins were examined for internal stop codons (excluding terminal stops), frameshift signals inferred from alignment operations, and truncation relative to the reference sequence. Copies exhibiting disabling mutations were classified as *putatively pseudogenized* (P). Copies with incomplete sequence recovery or ambiguous annotation were classified as *ambiguous* (A). Remaining copies without structural defects were classified as structurally intact (U0).

**Step 3: Functional divergence assessment**. Structurally intact copies were further evaluated for functional divergence using protein-level comparisons. Divergence from the reference gene used in the respective BLAST search (Step 1) was quantified using percent identity, protein length ratio, and relative divergence within each gene. Copies were classified as *putatively functionally changed* (F) if they met at least one of the following criteria: (i) sequence identity

< 90%; (ii) protein length ratio < 0.9 or > 1.1; or (iii) divergence exceeding a robust outlier threshold within each gene, a median absolute deviation (MAD)-based z-score (*z*_MAD_ > 3), where divergence (*d*) is calculated as *d* = 100 − identity(%). Copies not meeting these criteria were classified as *apparently unchanged* (U).

Final classifications comprised four categories: pseudogenized (P), functionally changed (F), unchanged (U), and ambiguous (A). This framework provides a mechanistic partitioning of CNV outcomes into loss-of-function, retention, and potential functional innovation, enabling direct comparison of evolutionary trajectories across gene families while accounting for both structural integrity and relative divergence.

### 2.6 Copy number variation-environment (CNV–ENV) association analysis

Environmental association analyses were conducted for both species across all rapidly-evolving gene families. Gene families were filtered prior to environmental modeling to remove families with little or no informative variation. For each gene family, we calculated the number of individuals, mean copy number, standard deviation, number of distinct copy-number states, number of non-zero observations, zero fraction, and minimum/maximum copy number. A gene family was retained if it satisfied all of the following criteria: *sd >* 0, at least five individuals with non-zero copy number, at least two distinct copy-number states, and a zero fraction ≤ 0.95. Families failing any of these criteria were excluded from downstream CNV–environment analyses.

To account for variation arising from neutral population structure, we incorporated population structure as covariates in all analyses using the first two principal components (PC1 and PC2) derived from independent SNP-based neutral datasets (i.e., intergenic regions, introns, and four-fold degenerate sites). For *A. thaliana*, SNP panels were generated from long-read sequence data for all accessions included in the CNV analysis. For *A. lyrata*, we used a published whole-genome SNP dataset (Scott et al., 2025). In both cases, the SNP datasets correspond to the same populations represented by the long-read assemblies analyzed here. Details of SNP panel construction and population structure inference are provided in Supplementary Information S4.

#### Technical and TE confounders

Before testing environmental associations, we evaluated whether global CNV structure was influenced by technical assembly-quality variables or genome-wide TE content. Three assembly-level technical metrics with low pairwise collinearity (Spearman |*ρ*| < 0.8) and one TE metric were retained: BUSCO completeness, contig N50, total assembly length (Mbp), and TE interspersed fraction (*TE*_*if*_).

Analyses were restricted to gene families showing within-dataset variation (*n*_unique_ > 1, *sd >* 0), and copy number was standardized within family to obtain *z*-scores (`cnv z`). Multivariate effects were assessed using partial redundancy analysis (partial RDA) with the standardized CNV matrix as the response, technical covariates or *TE*_*if*_ as explanatory variables, and PC1 and PC2 as conditioning variables: **CNV** ~ *covariate* + *Condition*(*PC1* + *PC2*). Significance of models and terms was tested using permutation tests (999 permutations).

In parallel, family-wise linear models (**cnv**_*z*_ ~ *covariate* + *PC1* + *PC2*) were fitted and corrected for multiple testing using the Benjamini–Hochberg procedure. Gene families showing significant associations were flagged as potentially confounded.

#### 2.6.1 Environmental predictors

Bioclimatic variables (Bio1–Bio19) were obtained from the CHELSA climatic database (Karger et al., 2017) at 30 arc sec resolution and edaphic variables were downloaded from the SoilGrids database (Poggio et al., 2021). We used soil pH, cation exchange capacity (CEC), and soil organic carbon (SOC) at depths 0-30 cm (Tab. S5). All environmental variables were screened in several steps to reduce multicollinearity. First, predictors with near-zero variance were identified and removed. Then pairwise Spearman correlations were calculated among retained predictors and highly correlated variables (|*ρ*| > 0.8) were removed. Finally, variance inflation factors (VIFs) were calculated iteratively, and variables with VIF > 6 were removed one at a time until all retained predictors satisfied the threshold.

#### 2.6.2 Partial redundancy analyses of CNV–environment associations

To examine multivariate associations between CNV and environmental gradients, we constructed a response matrix from the filtered gene-family copy numbers, containing family-wise standardized copy numbers (**CNV**_*z*_), obtained by *z*-transforming copy number within each gene family across assemblies. This standardization removes scale differences among gene families and ensures that highly variable or high-copy families do not disproportionately influence multivariate analyses. Partial RDAs were fitted using the `vegan` package. The main models tested the joint effect of climate and soil variables while conditioning on population structure: **CNV**_*z*_ ~ climate + soil + *Condition*(*PC1* + *PC2*). Additionally, we also used ploidy as a conditional variable in *A. lyrata* to account for ploidy differences.

To partition the unique contributions of climate and soil, two additional sets of partial RDAs were fitted. Pure climate effects were estimated by conditioning on both soil variables and structure: **CNV** ~ climate + *Condition*(soil + *PC1* + *PC2*); Pure soil effects were estimated analogously by conditioning on climate variables and structure: **CNV** ~ soil + *Condition*(climate + *PC1* + *PC2*). For each model, overall significance and term-wise significance were evaluated using permutation tests (999 permutations).

##### Variance partitioning

To quantify the unique and shared contributions of climate, soil, and neutral structure to CNV variation, we performed variance partitioning on both raw and family-standardized CNV matrices. Three explanatory sets were specified: climate variables, soil variables, and the neutral structure covariates (PC1 and PC2). This analysis decomposed the explained variation into unique fractions attributable to each predictor set and their overlaps.

#### 2.6.3 Family-wise CNV–environment association tests

In addition to the multivariate analyses, we carried out family-wise univariate association testing using family-standardized copy number (*CNV*_*z*_) as the response. For each gene family and each environmental predictor, we fitted linear models of the form: **cnv**_*z*_ ~ environmental predictor+ (*PC1* + *PC2*). Same as in partial RDA, we also used ploidy as a conditional variable in *A. lyrata* to account for ploidy differences.

Predictor variables were standardized prior to modeling so that regression coefficients were directly comparable among variables. For each environmental predictor, *P*-values across families were corrected using the Benjamini–Hochberg FDR procedure. Summary statistics included the number of families significant at nominal *P <* 0.05, the number significant at FDR < 0.05, median absolute effect size, and median adjusted *R*^2^. These family-wise analyses complemented the partial RDA by identifying specific gene families whose CNV profiles were associated with individual environmental gradients after accounting for population structure.

## 3 Results

### 3.1 Comprehensive Identification of Gene Families

Our gene family assignment using OrthoFinder2 yielded a well-resolved, rooted species tree with strong bootstrap support (Fig. 1), consistent with established phylogenies of the order *Brassicales* (e.g., Hendriks et al., 2023). Across all five species, OrthoFinder2 annotated a total of 136,809 genes, of which 92% were assigned to 22,083 orthogroups. Among the ingroup taxa, over 93% of genes were clustered into orthogroups, whereas the outgroup had a lower assignment rate of 79% (Table 1). *Arabidopsis halleri* showed the highest proportion of genes in orthogroups (97.5%), while *A. alpina* exhibited the greatest number of species-specific orthogroups (292, comprising 10% of all its genes). In contrast, *A. thaliana* had the fewest species-specific orthogroups (99, 1.2%).

**Table 1:** Summary statistics of gene content and orthogroup (OG) assignment across five species used in the OrthoFinder2 analysis. The table reports the total number of predicted genes, the proportion assigned to orthogroups (shared gene families), and the number and percentage of unassigned genes for each species. It also shows how many orthogroups each species contributes to (number of orthogroups containing at least one gene from that species– “OGs represented in species”) as well as the number and proportion of species-specific orthogroups (i.e., orthogroups unique to a single species).

|  | *A. alpina* | *A. halleri* | *A. lyrata* | *A. thaliana* | *C. papaya* |
| --- | --- | --- | --- | --- | --- |
| **Ecotype** | Pajares | Lan3.1 | MN47 | Col-0 | — |
| **Annotation** | V4 | v2.1.0 | v2.1 | Araport11 | ASGPBv0.4 |
| **BUSCO (%)** | 79.8 | 94.0 | 98.2 | 99.3 | 72.0 |
| **N50 (Mb)** | 0.79 | 23.0 | 24.5 | 23.5 | 1.18 |
| **Genes** | 21,609 | 28,722 | 31,073 | 27,654 | 27,751 |
| **Genes in OGs** | 20,144 (93.2%) | 27,999 (97.5%) | 29,383 (94.6%) | 26,442 (95.6%) | 21,896 (78.9%) |
| **Unassigned genes** | 1,465 (6.8%) | 723 (2.5%) | 1,690 (5.4%) | 1,212 (4.4%) | 5,855 (21.1%) |
| **OGs represented in species** | 13,443 (60.9%) | 18,734 (84.8%) | 19,735 (89.4%) | 19,228 (87.1%) | 13,315 (60.3%) |
| **Species-specific OGs** | 292 | 241 | 321 | 99 | 889 |
| **Genes in species-specific OGs** | 2,162 (10.0%) | 1,505 (5.2%) | 992 (3.2%) | 341 (1.2%) | 4,635 (16.7%) |

**Table 2:** Summary of gene family copy number variation within and between *A. lyrata* and *A. thaliana*. “Variable families” denotes the proportion of gene families with non-zero variance in copy number across assemblies. “Copy range” indicates the median copy number per family with the full range across all families. “Variance” and “CV” (coefficient of variation) are calculated per gene family across assemblies and summarized as median [range]. “Dispersion” represents the variance-to-mean ratio of copy number per family, summarizing dispersion in CNV across assemblies. “CNV prevalence *P* “ corresponds to an exact binomial test assessing whether the proportion of variable families is significantly greater than 0.5 within each species.

| Species | Families | Variable families | Copy range | Variance | CV | Dispersion | CNV prevalence $P$ |
| --- | --- | --- | --- | --- | --- | --- | --- |
| *A. lyrata* | 224 | 0.882 | 2.5 [0–175] | 0.548<br>[0–2993.029] | 0.158<br>[0–1.075] | 0.140<br>[0–17.518] | $2.28 \times 10^{-34}$ |
| *A. thaliana* | 228 | 0.689 | 1.0 [0–103] | 0.057<br>[0–328.267] | 0.060<br>[0–2.255] | 0.015<br>[0–8.118] | $6.14 \times 10^{-9}$ |

**Figure 1:**
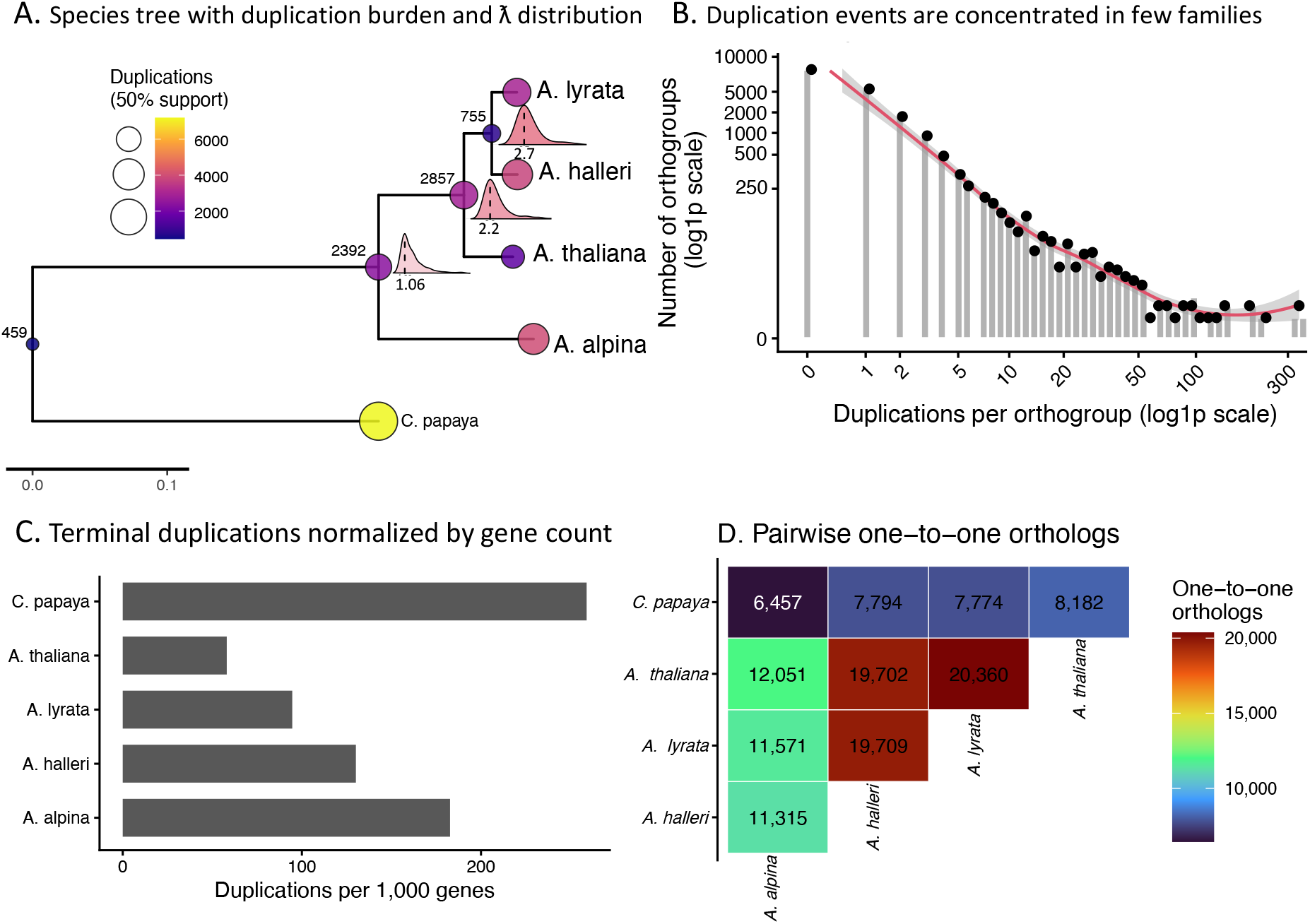
Comparative genomic dynamics of gene families across Brassicaceae species. **(A)** Time-calibrated species tree with inferred gene duplication burden and gene family evolutionary rates. Circles at internal nodes indicate the number of gene duplication events (≥50% support), with size and color proportional to duplication counts; values in black denote the absolute number of duplications at each node. Density plots at nodes represent the distribution of gene family evolutionary rates (*λ*) inferred from CAFE5, with dashed lines indicating mean *λ* and color intensity reflecting its magnitude. **(B)** Distribution of duplication events per orthogroup (log_1*p*_ scale), showing that duplications are highly uneven and concentrated in a small subset of gene families. **(C)** Terminal duplication rates normalized per 1,000 genes across species, enabling comparison of lineage-specific duplication intensity independent of genome size. **(D)** Pairwise counts of one-to-one orthologs among species, illustrating overall genomic similarity, with the highest values observed between *A. thaliana* and *A. lyrata*. Together, these results reveal pronounced heterogeneity in gene family evolution, with lineage–specific differences in duplication burden, evolutionary rates, and gene family turnover across Brassicaceae species.

Within in-group species, a large proportion of genes (46-66%) were found in single-copy orthogroups, with *A. thaliana* having the highest percentage (66.7%). *A. halleri* showed the greatest number of large gene families (129 orthogroups with 16 or more genes), while *A. thaliana* had the fewest (48 orthogroups). Orthology patterns also support close relationships among the ingroup species. The highest one-to-one ortholog counts were observed between *A. lyrata* and *A. thaliana* (20,360) and between *A. halleri* and both *A. lyrata* (19,709) and *A. thaliana* (19,702). Ortholog counts involving *C. papaya* were much lower, consistent with its role as an outgroup (Fig. 1D).

Gene duplication was widespread but uneven among lineages (Fig. 1A-C). Terminal duplications were highest in *A. alpina* (3,948) and *A. halleri* (3,738), followed by *A. lyrata* (2,940), whereas *A. thaliana* showed fewer terminal duplications (1,605). This supports the broader pattern that duplicated gene-family dynamics are reduced or more constrained in *A. thaliana* relative to the other Brassicaceae species. Across orthogroups, 8,646 of 17,017 orthogroups with inferred duplications had at least one duplication event showing a strongly skewed distribution: the median was 1 duplication per orthogroup, but the maximum reached 360, indicating that a minority of large, rapidly evolving families account for a disproportionate share of duplication events (Fig. 1B).

### 3.2 Modeling Gene Family Evolution and Rapidly Evolving Families

We analyzed the evolutionary dynamics of the 22,083 gene families using CAFE 5, which models gene family size changes under a birth-and-death process. The input included the gene copy number per species and the dated, rooted species tree. The species tree from OrthoFinder was calibrated using the `makeChronosCalib` function from the `APE` R package (Paradis and Schliep, 2019), based on a divergence estimate of 4-6 Mya between *A. thaliana* and the rest of the *Arabidopsis* species (Novikova et al., 2016; Hohmann et al., 2015).

To identify the most suitable evolutionary rate model, we compared models with one to five gamma distribution of evolutionary rate classes (*γ* = 1–5). Both Akaike (AIC) / Bayesian (BIC) information criteria and log-likelihood supported the base model (*γ* = 1) as the best fit (Fig. S1). The estimated average gene family turnover rate under this model was *λ* = 0.0118, with the maximum rate for this topology being *λ*_max_ = 0.0238. Incorporating an error model to account for annotation errors did not significantly alter the estimated rate (*λ* = 0.0109, [error rate] *ε* = 0.0232), confirming the robustness of the base model. We therefore retained the base model for downstream analyses.

Based on the base model, CAFE 5 identified 1,592 gene families with significant expansion or contraction (p<0.05). For a more stringent and biologically relevant subset, we applied a significance threshold of p<0.01, resulting in 821 gene families classified as “rapidly-evolving.” To facilitate comparative and functional analyses, we filtered this list further by retaining only the gene families present in at least three species and containing at least one gene from *A. thaliana*, enabling downstream annotation and functional prediction. This yielded a final set of 231 rapidly-evolving gene families for in-depth exploration.

### 3.3 Functional Categories of Fast-Evolving Gene Families

Based on *A. thaliana* gene ontology (GO) predictions, the 231 significantly fast-evolving gene families were classified into 10 major functional categories (Fig. 2). The largest proportion comprised **multi-function proteins** (21.6%), followed by **genetic information & regulation** (21.2%). **Defense and stress response genes** accounted for 17.7%, and 15.2% were uncategorized.

**Figure 2:**
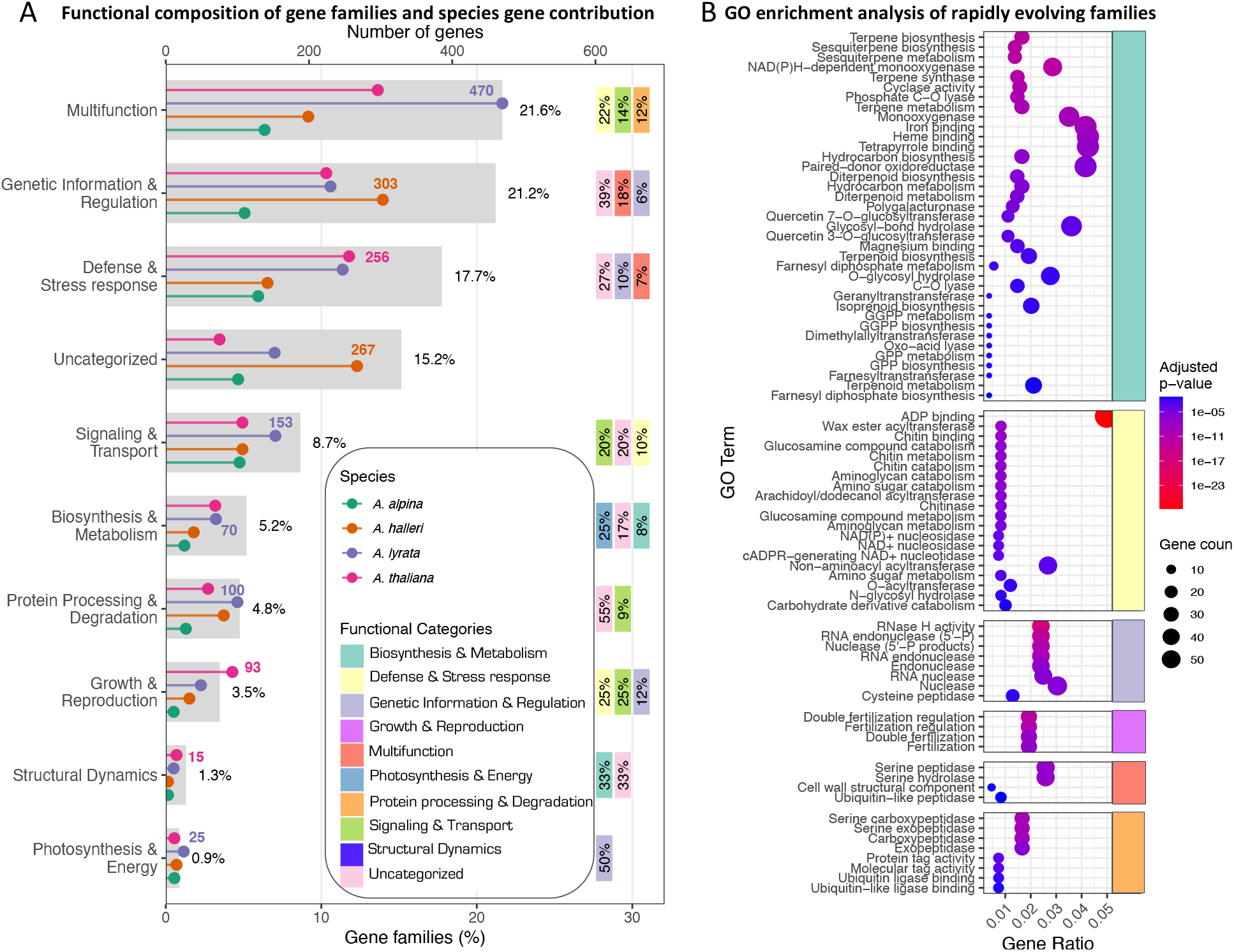
Functional classification and GO enrichment of rapidly evolving gene families. **(A)** Functional composition of rapidly evolving gene families. Grey bars indicate the proportion of gene families assigned to each primary functional category, while lollipop markers show the number of genes contributed by each species within those categories. Functional categories are defined based on the dominant annotated function of at least one gene per orthogroup. Colored tiles to the right represent secondary functional categories within each primary category, with percentages indicating their relative contribution. Functional annotations were derived from *A. thaliana* gene models using data from TAIR and UniProt. **(B)** Gene Ontology (GO) enrichment analysis of rapidly evolving gene families. Only significantly enriched GO terms are shown (Benjamini–Hochberg adjusted *p*-value). Gene ratio represents the proportion of genes associated with a given GO term relative to all genes included in the analysis. GO terms were simplified for visualization; the complete list of terms and enrichment statistics is provided in Tab. S6. Category colors are same as in A.

Notably, many gene families contained members with diverse gene ontology terms. In such cases, we assigned the family to the most frequently occurring functional category among its members. A comprehensive list of gene families, their constituent genes, and assigned functions is provided in Tab. S11. Further, a GO enrichment analysis indicated that both “Defense and Stress-Response” and “Biosynthesis and Metabolism” related gene families were significantly enriched with former having the highest enrichment (gene ratio: 0.05; adjusted p-value: 1.8 ×10^−29^: see Fig 2B and Tab. S6).

### 3.4 Gene family composition and chromosomal organization across species

Our analysis of gene family composition revealed that more than 50% (124) of gene families exhibited functional diversification, defined as the presence of gene family members encoding functionally distinct proteins. Such diversification is commonly associated with neofunctionalization or subfunctionalization following gene duplication, facilitating the evolution of novel biological roles or ecological adaptations (Innan and Kondrashov, 2010). In contrast, functionally conserved families—comprising members with similar molecular and regulatory functions—likely evolve under strong purifying selection, as typically observed for essential or dosage-sensitive genes (e.g., Ohno, 1970; Birchler and Veitia, 2012).

Chromosomal clustering of gene family members (i.e., localization on the same chromosome versus dispersed across chromosomes) differed significantly among the four Brassicaceae species (*χ*^2^ = 15.176, *df* = 3, *P* = 0.0017) (Tab. S1A), indicating lineage-specific patterns of genome organization. Notably, *A. lyrata* exhibited significantly fewer clustered gene families than the other species (*P <* 0.006; Tab. S1B), consistent with a greater degree of inter-chromosomal gene dispersal. Such differences likely reflect distinct evolutionary mechanisms underlying gene family expansion: clustering is typically associated with tandem or local duplication, whereas dispersed distributions may result from translocations, older duplication events, or transposable element (TE)-mediated processes. The reduced clustering in *A. lyrata* therefore suggests a larger contribution of genome rearrangements to its gene family evolution.

### 3.5 Transposable element enrichment in duplicated gene families

Across *A. thaliana* and *A. lyrata*, duplicated-family genes consistently exhibited higher TE content than single-copy genes across all spatial scales, from gene bodies to extended flanking regions (Fig. 3A-C). TE fraction increased with window size, and differences between duplicated-family and single-copy genes became more pronounced at broader genomic scales, indicating that duplicated gene families are preferentially located within TE-rich genomic environments. TE occupancy showed substantial heterogeneity among loci. Duplicated-family genes displayed broader and more right-skewed distributions of TE fraction, with a subset of loci occurring in highly TE-dense regions (Fig. 3C). This pattern was more pronounced in *A. lyrata*, where both TE fraction and variability were higher, consistent with its more TE-rich genome and in line with the genome size reduction observed in *A. thaliana* (de la Chaux et al., 2012).

**Figure 3:**
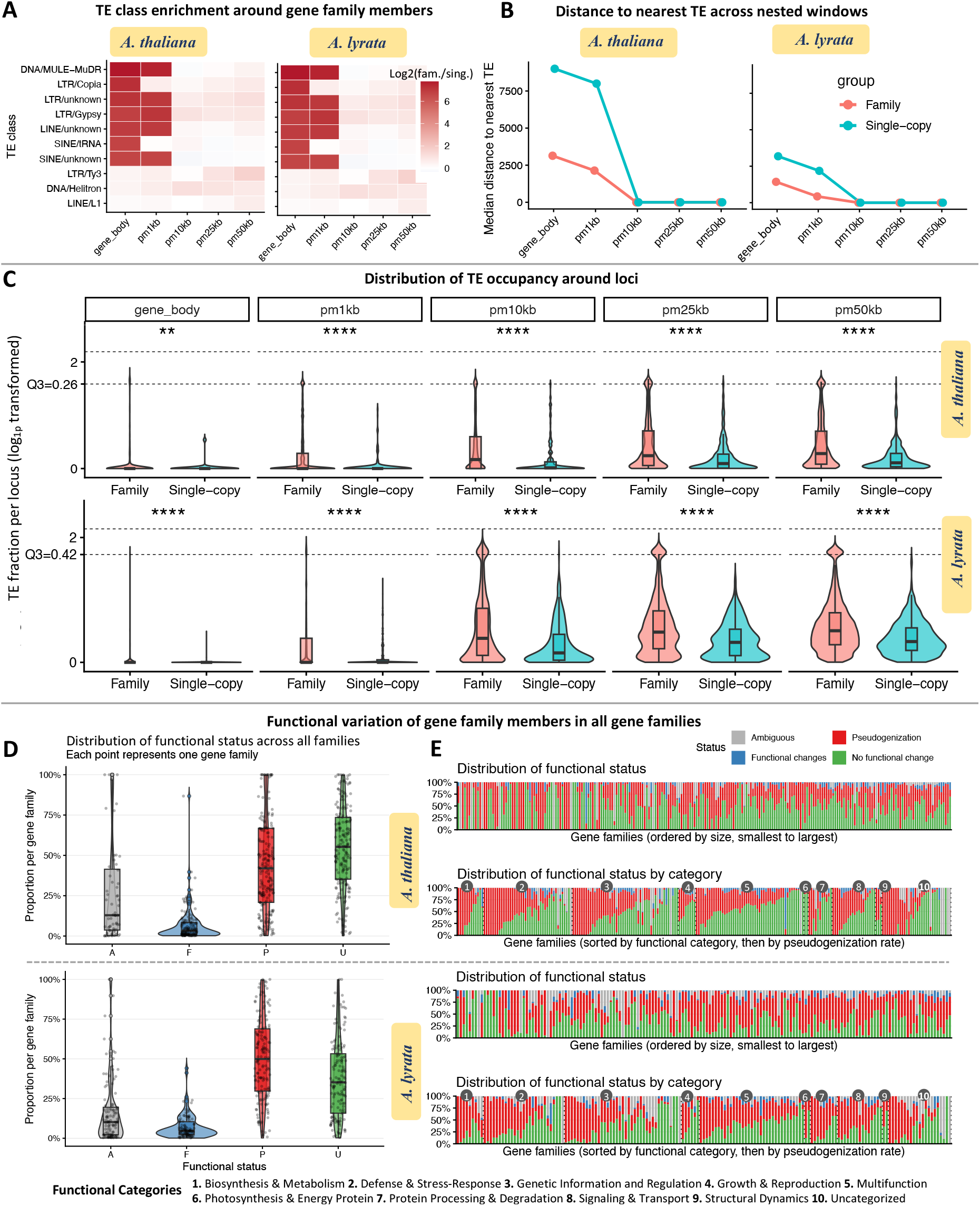
Transposable element (TE) association and functional fate of duplicated gene family members. **A–C:** Association of gene family members with transposable elements compared to single-copy genes. **A:** Enrichment of TE classes around duplicated gene family members relative to single-copy genes across nested genomic windows (gene body, ±1 kb, ±10 kb, ±25 kb, ±50 kb), shown as log_2_ fold-change. **B:** Median distance to the nearest TE for gene family members and single-copy genes across the same genomic windows, illustrating spatial proximity to TEs. **C:** Distribution of TE fraction per locus (log_1*p*_-transformed) for gene family members versus single-copy genes, with significance levels indicated (asterisks); plots are truncated at Q3 to improve visualization. **D–E:** Functional fate of duplicated gene family members across all gene families. **D:** Distribution of functional status per gene family, proportions correspond to genes classified as no functional change (U), functional change (F), pseudogenization (P), or ambiguous (A). **E:** Composition of functional outcomes across gene families, shown as families and proportion of genes in color bars. Results are shown for *A. thaliana* and *A. lyrata*.

TE class-specific analyses showed that enrichment was driven by similar TE classes in both species, particularly LTR retrotransposons (LTR/Gypsy, LTR/Copia) and DNA transposons (DNA/MULE-MuDR, DNA/Helitron), while SINE elements showed weak enrichment (Fig. 3A). Enrichment increased with genomic window size and was consistently stronger in *A. lyrata*.

All differences between duplicated-family and single-copy genes were statistically significant across genomic windows (Wilcoxon tests, *p* ≪ 0.001; Supplementary Tab. S8). Together, these results indicate that TE association with duplicated gene families is conserved across species but markedly stronger in *A. lyrata*, suggesting a greater role of TE-rich genomic environments in shaping gene family evolution in genomes with higher TE content.

#### 3.5.1 Functional and evolutionary fate of duplicated gene families

We quantified the structural and functional status of gene copies across all rapidly evolving gene families in *A. thaliana* and *A. lyrata*. A direct comparison between the two species reveals two contrasting regimes in the evolutionary fate of duplicated gene copies (Fig. 3D–E; Tab. S7.1–.2). In *A. thaliana*, duplicated copies are more frequently retained in a structurally intact state (U0~46.7%) than lost through pseudogenization (P~41.9%), and half of all gene families (U~50.0%) are dominated by functionally conserved copies. In contrast, *A. lyrata* shows a clear shift toward gene loss, with pseudogenization exceeding retention at both the copy level (P~49.0% vs. U~35.3%) and the family level (P~54.5% vs. U~34.5%). Thus, while both species exhibit substantial turnover following duplication, the balance between retention and loss is markedly different.

Despite these differences, both species share a common feature: functional divergence (F) is relatively rare in absolute terms (5.6% in *A. thaliana*; 7.5% in *A. lyrata*) and seldom dominates entire gene families (≤1.2%). Instead, functionally divergent copies typically occur alongside intact copies within the same family, indicating that diversification arises in a subset of duplicates rather than through wholesale functional shifts at the family level. This suggests that neofunctionalization or subfunctionalization operates opportunistically within retained copies, rather than being the primary outcome of gene duplication.

The contrasting dominance of functional states across families highlights distinct evolutionary trajectories. In *A. thaliana*, the coexistence of high retention (50.0% of families) and substantial but secondary pseudogenization (43.0%) reflects a relatively balanced birth–death dynamic, where duplicated genes are frequently maintained under purifying selection while a considerable fraction is lost. In *A. lyrata*, however, the predominance of pseudogenized families (54.5%) and reduced retention indicates a birth–death process skewed toward gene loss, suggesting either weaker selective constraints on duplicate maintenance or higher rates of genome turnover. We did not find any significant association between gene family size or functional category and the prevalence of pseudogenization or functional divergence in either species (Fig. 3E), indicating that these evolutionary outcomes are largely independent of gene family composition or functional class.

Together, these patterns point to lineage-specific differences in how CNV contributes to genome evolution. *A. thaliana* appears to favor retention of duplicated genes, maintaining a larger pool of functional copies that can potentially contribute to phenotypic flexibility, even though functional divergence remains limited. In contrast, *A. lyrata* exhibits a more transient duplication landscape, where many duplicated copies are rapidly degraded, leading to fewer retained functional copies but similar, low levels of diversification among those that persist. These results indicate that closely related species can differ fundamentally in the balance between retention, loss, and innovation following gene duplication, with important implications for their evolutionary potential and genomic plasticity.

### 3.6 Copy number variation across rapidly evolving gene families

#### 3.6.1 Global patterns of copy-number variation

From the initial 231 rapidly evolving gene families, three organellar (plastid or mitochondria) families were removed due to absence of data in the long-read assemblies–and four families were absent in *A. lyrata*. Across the remaining gene families, copy number varied substantially both within and between species. Total gene family copy counts per assembly were consistently higher in *A. lyrata* than in *A. thaliana*, indicating a general expansion of gene family sizes in *A. lyrata* (Fig.S2; Tab. 2&S2). A generalized linear mixed model (negative binomial) confirmed a strong species effect on copy number (likelihood ratio test: *χ*^2^ = 464.7, df = 1, *P <* 2.2 × 10^−16^), demonstrating genome-wide differences beyond family-specific variation. Consistently, multivariate analyses showed clear separation between species, with species identity explaining most of the variation in CNV structure (PERMANOVA: *R*^2^ = 0.863, *P* = 0.001).

Within species, CNV was highly heterogeneous across gene families. In *A. thaliana*, many families (~31%) showed little or no variation, indicating largely stable copy numbers. In contrast, *A. lyrata* exhibited widespread variation, with higher variance and coefficients of variation across families, reflecting substantial heterogeneity among individuals (Fig. S3A&B).

At the family level, between-species differences were pervasive. A large proportion of families showed significant copy-number differences after multiple-testing correction (FDR-adjusted Wilcoxon tests, *P* ≪ 0.001), with a general shift toward higher copy numbers in *A. lyrata*. Notably, several families invariant in *A. thaliana* were variable in *A. lyrata*, highlighting lineage-specific CNV dynamics as a key driver of species divergence.

#### 3.6.2 Geographic structuring of CNV profiles

Principal component analysis of combined CNV profiles revealed a striking separation between *A. thaliana* and *A. lyrata*, with the first principal component explaining the vast majority of variation (PC1 = 87%; Fig. S5). Assemblies from the two species formed completely non-overlapping clusters along this axis, indicating that interspecific differences in gene family copy number dominate the global CNV landscape, consistent with their evolutionary divergence. In contrast, the second principal component explained only a minor fraction of variation (PC2 = 1.2%) and primarily captured within-species structure, indicating that CNV divergence between species is driven by coordinated shifts in gene family copy numbers rather than subtle multivariate differences.

Within species, CNV profiles exhibited distinct but contrasting patterns. In *A. thaliana*, clustering was structured but partially overlapping and broadly consistent with known geographic and demographic patterns for the species (Fig. 4B). European accessions formed a dense central cluster, reflecting relatively homogeneous CNV profiles across the core range. Southern populations, including Iberian, North African, and Cape Verde accessions, largely grouped together, consistent with shared CNV signatures associated with relict lineages, while Madeira accessions were more distinctly separated, indicating stronger divergence likely driven by prolonged isolation. Asian accessions formed a partially distinct cluster with overlap into eastern European and Central Asian populations, suggesting gradual differentiation rather than discrete separation. These patterns are consistent with previous SNP- and CNV-based studies (e.g. Alonso-Blanco et al., 2016; Göktay et al., 2020; Kang et al., 2023), but the long-read assembly–based approach used here reveals a more continuous structure, suggesting that CNV variation transitions more gradually across geographic space than previously inferred (e.g., Lian et al., 2024).

**Figure 4:**
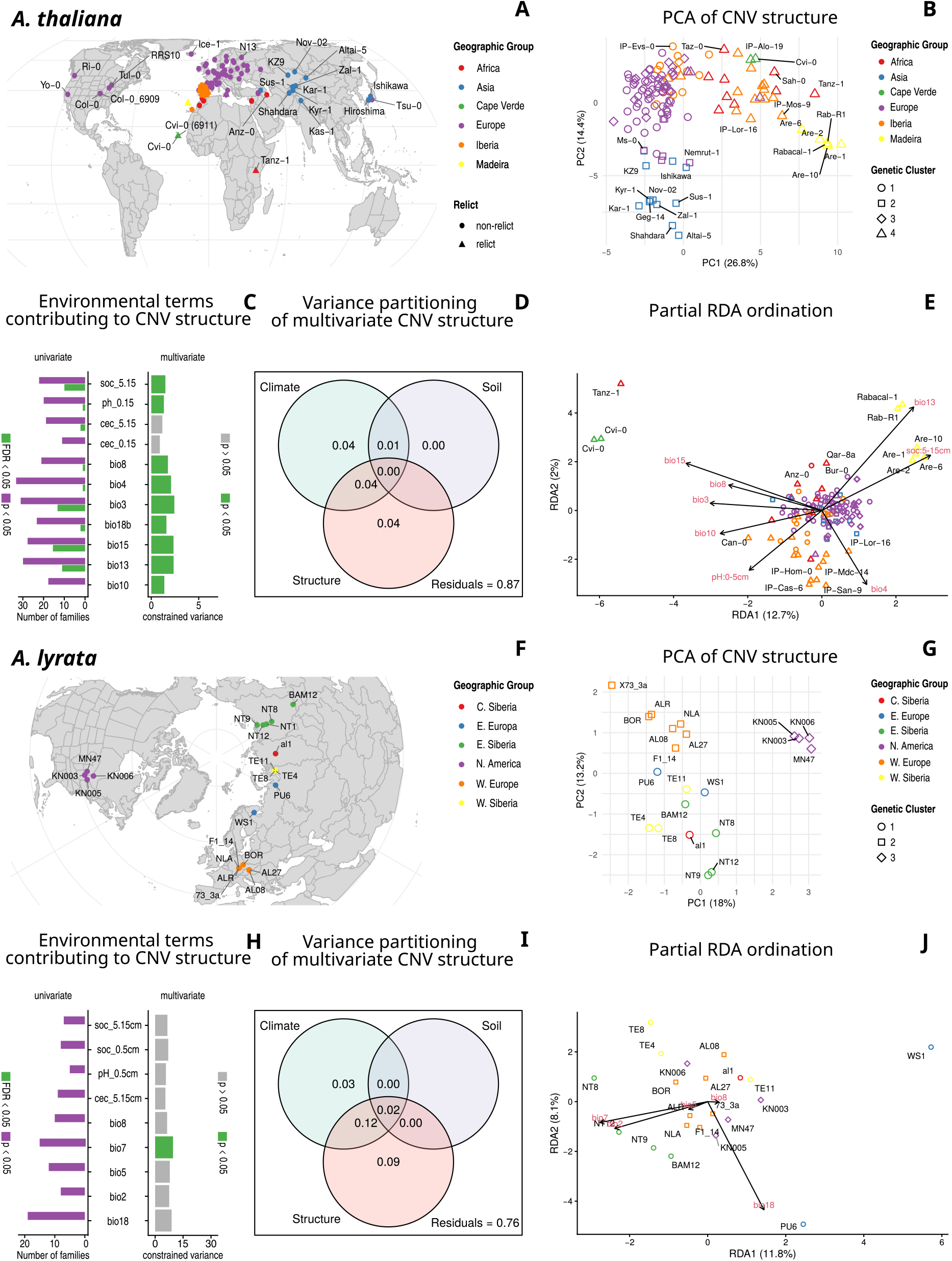
Geographic structuring of CNV in rapidly evolving gene families. **A & F:** Sampling locations of *A. thaliana* and *A. lyrata*, respectively. Colors indicate geographic clusters based on previous studies (Wlodzimierz et al., 2023; Alonso-Blanco et al., 2016; Scott et al., 2025). In *A. thaliana*, point shapes additionally denote relict status (Alonso-Blanco et al., 2016). Map boundaries follow TDWG level III floristic regions (Brummitt et al., 2001). **B & G:** PCA of CNV profiles across gene families. Colors correspond to geographic clusters (A, F), and shapes represent genetic clusters inferred from hierarchical clustering of covariance matrices derived from putatively neutral loci (four-fold degenerate sites, introns, and intergenic regions; Tables S10.1–S10.2). **C & H:** Contribution and significance of environmental predictors to CNV variation in multivariate (right; RDA) and univariate (left) analyses. **D & I:** Variance partitioning of CNV among climate, soil, and population structure. **E & J:** Partial RDA ordination of environmental effects on CNV (*cnv* ~ *climate* + *soil* | *structure*). Arrows indicate the direction and strength of environmental associations. Bioclimatic variables are derived from CHELSA (Karger et al., 2017) and soil variables from SoilGrids (Poggio et al., 2021) (see Table S5 for details).

In contrast, *A. lyrata* exhibited stronger and more discrete multivariate structure (Fig. 4G). Assemblies clustered into well-defined continental groups, with North American populations clearly separated from Eurasian lineages along the primary axis, and additional structure distinguishing eastern Siberian from European and western Siberian populations. Compared to *A. thaliana*, clustering was tighter and less overlap, indicating that CNV variation in *A. lyrata* is driven by fewer, larger-effect gene family changes aligned with major phylogeographic divisions. These patterns are consistent with SNP-based studies (Scott et al., 2025; Hämälä et al., 2018b) but are further resolved by CNV, which captures structural genomic variation shaped by demographic history and local adaptation. This is the first account of CNV structure distribution within the species.

Taken together, these results reveal a hierarchical structure of CNV variation across *Arabidopsis*, with species identity dominating global differentiation and contrasting within-species architectures: a diffuse and continuous CNV landscape in *A. thaliana* versus a more discrete and strongly structured pattern in *A. lyrata*. These differences are consistent with stronger gene family expansion dynamics and greater structural genomic heterogeneity in *A. lyrata*.

### 3.7 Association between CNV and environmental variation

Preliminary analyses indicated minimal confounding effects of assembly quality and genome-wide TE content, as neither factor showed consistent multivariate or family-wise associations with CNV (Tab. S9). After removing non-variable families, 157 and 201 gene families were retained for *A. thaliana* and *A. lyrata*, respectively for the CNV-ENV analysis.

Variance partitioning and partial RDA revealed that environmentally associated CNV is present in both species but differs in structure and interpretability. In *A. thaliana*, the full model explained 21.2% of total variance (adjusted *R*^2^ = 0.126), with substantial residual variation. Climate and population structure contributed comparable independent fractions (climate: 0.036; structure: 0.038), while soil effects were small but positive (0.005). Shared fractions, particularly between climate and structure (0.038), were moderate, indicating partial collinearity but also a distinct climatic signal (Fig. 4D; Tab. S5). In contrast, *A. lyrata* showed a higher total variance explained (adjusted *R*^2^ = 0.224), but with a different partitioning pattern: population structure accounted for the largest independent component (0.092), whereas the unique climatic contribution was smaller (0.034) and soil effects were negligible after adjustment. A substantial proportion of variance was shared among predictors, especially between climate and structure (0.075), indicating that environmental associations are largely aligned with phylogeographic structure (Fig. 4I).

Ordination of partial RDA models was consistent with these patterns. In *A. thaliana*, most populations remained tightly clustered (e.g., European populations) after conditioning on structure, reflecting broadly similar CNV profiles and limited environmentally associated differentiation among these samples rather than an absence of overall variation. Furthermore, climatically distinct populations (e.g., Cape Verde, Madeira, and Tanzania) were displaced along environmental gradients, suggesting localized CNV responses. In *A. lyrata*, conditioning on population structure more strongly altered population relationships in the ordination, yet the residual CNV variation remained predominantly structured along phylogeographic rather than environmental axes. Albeit with limited separation independent of structure, all Siberian populations showed patterns of climatic gradient-associated clustering (Fig. 4E, J).

Family-wise analyses further emphasized these contrasting architectures. In *A. thaliana*, CNV–environment associations were distributed across multiple gene families, with several climatic variables (notably Bio3: *Isothermality*, Bio13: *Precipitation of the wettest month*, and Bio15: *Precipitation seasonality*) yielding multiple FDR-significant associations (Fig. 4C). Although per-family effect sizes were modest, signals were consistent across functionally relevant categories, including defense and stress response, signaling, and genetic regulation, and several families showed associations with multiple environmental predictors, indicating a diffuse, polygenic pattern. In *A. lyrata*, associations were more limited after multiple-testing correction. Nevertheless, 81 gene families showed nominal associations—primarily with *Temperature annual range* (Bio7) and *Precipitation of the warmest quarter* (Bio18)—providing informative signals given the limited statistical power imposed by the small number of populations (n = 24) relative to the number of tests. Although none retained FDR-significance, these nominal associations spanned diverse functional categories and likely capture biologically relevant but subtle or population-specific CNV–environment relationships.

Functional enrichment analysis refined these patterns. In *A. thaliana*, significantly associated families were strongly enriched for defense- and stress-response functions (odds ratio = 4, *p <<* 0.001), indicating that environmentally associated CNV is biased toward ecologically relevant pathways. In *A. lyrata*, enrichment of structural and genomic dynamics-related families was detectable but limited to a relatively small subset of significant families. Instead, multifunctional and defense- and stress-response gene families accounted for the largest number of significantly associated families (Fig. 5B-C), with multifunctional categories frequently encompassing defense-related roles as secondary annotations. This pattern suggests that, despite differences in enrichment profiles, environmentally associated CNV in both species is predominantly concentrated in defense-related functional space, highlighting a shared role of CNV in mediating stress and environmental responses.

**Figure 5:**
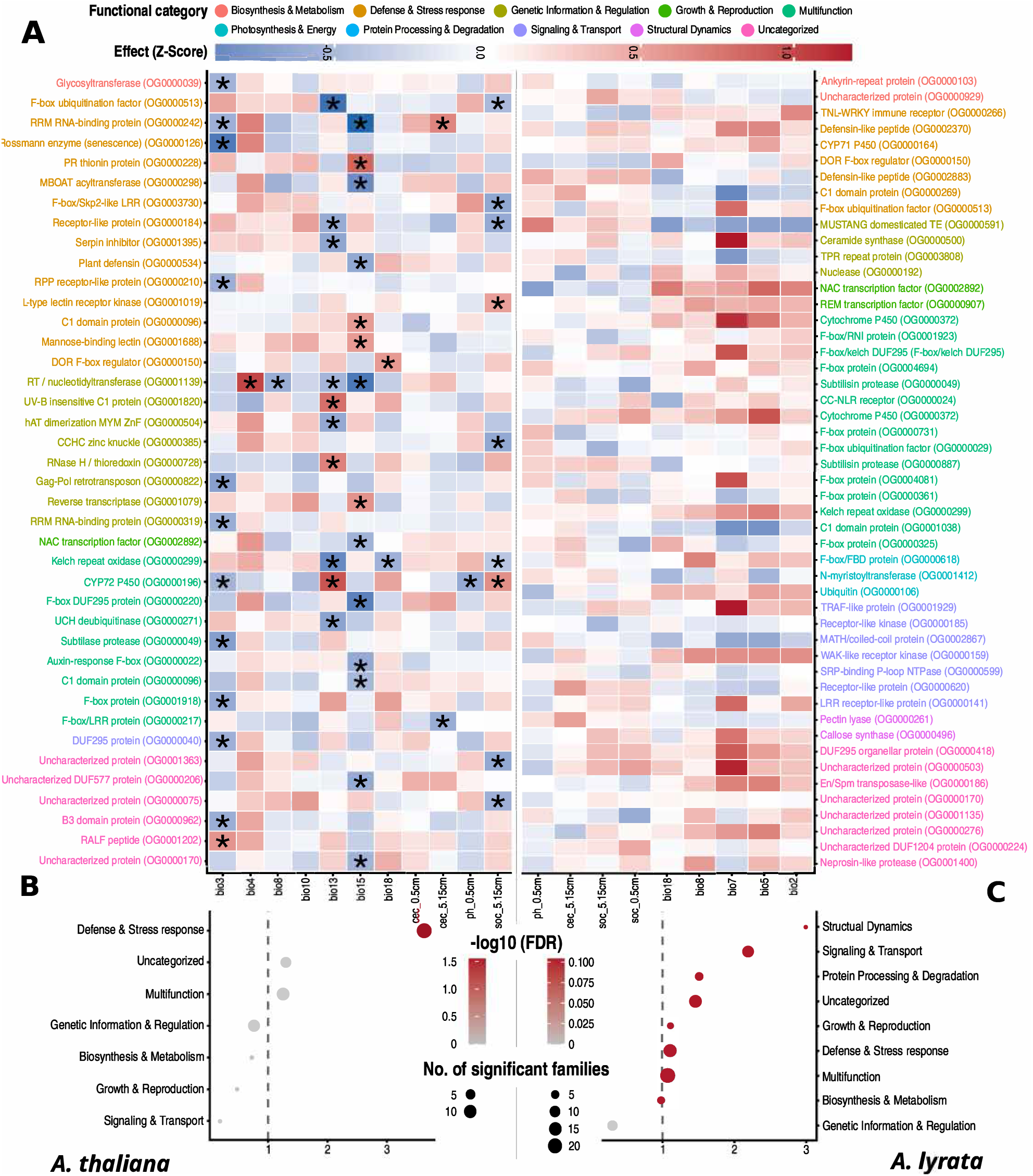
A: Gene family–level associations between copy number variation (CNV) and environmental variables. Heatmaps show univariate associations between gene family CNV (z-scores) and environmental predictors for gene families with significant CNV–environment relationships in *A. thaliana* (left) and *A. lyrata* (right). Each row represents a gene family and each column an environmental variable, including bioclimatic variables (bio2–bio18) and soil properties (e.g., cation exchange capacity, pH, and soil organic carbon at different depths). Color intensity indicates the standardized effect size (Z-score), with red representing positive and blue negative associations. Stars denote statistically significant associations after multiple-testing correction (FDR < 0.05), while all displayed families showed nominal significance (P < 0.05) in the univariate tests. Gene families are annotated by functional category, highlighting the diversity of biological processes involved. **B-C: Functional category enrichment of gene families showing CNV–environment associations**. Functional enrichment results from family-wise univariate analyses are shown for *A. thaliana* (B) and *A. lyrata* (C). Points represent functional categories, with enrichment expressed as odds ratios,

Taken together, these results indicate two contrasting regimes of CNV–environment association. In *A. thaliana*, CNV exhibits a modest but partially independent climatic signal distributed across many gene families, consistent with a diffuse, polygenic response. In *A. lyrata*, the overall signal is more strongly structured by population history, with environmental associations largely overlapping with phylogeographic patterns and concentrated in fewer gene families.

These findings place previous CNV studies in a broader comparative framework. Earlier work in *Arabidopsis* and other plant systems has largely focused on selected loci or candidate gene families, demonstrating roles of CNVs in adaptive and metabolic variation (Shirai et al., 2017; Wei et al., 2025; Wilson et al., 2025). In contrast, our genome-wide, multi-species analysis shows that these signals scale up to the level of gene-family landscapes, revealing consistent enrichment of defense-related functions while also exposing pronounced differences in the underlying architecture of CNV–environment associations between closely related species. Specifically, while *A. thaliana* exhibits a diffuse, polygenic CNV response to environmental gradients, *A. lyrata* shows patterns more tightly coupled to phylogeographic structure and concentrated in fewer gene families. This comparative perspective demonstrates that CNV–environment relationships are not only shaped by selection on specific loci, but emerge from the interaction between gene family evolution, demographic history, and genome organization.

## 4 Discussion

Our comparative analysis shows that gene family evolution in Brassicaceae is shaped by both conserved macroevolutionary processes and lineage-specific genomic dynamics. Most gene families fit a homogeneous birth–death model, consistent with gradual accumulation of duplication and loss over evolutionary time (Lynch and Conery, 2000; Hahn et al., 2005). However, duplication events were unevenly distributed, with a minority of rapidly evolving families accounting for much of the turnover. This pattern is consistent with broader observations in angiosperms (Cannon et al., 2004; Panchy et al., 2016), but highlights that genome-wide stability can coexist with strong family-specific evolutionary lability.

Rapidly evolving families were concentrated in defense, stress response, signaling, and metabolism-related functions. These included receptor-like proteins, lectin-like proteins, defensin/ thionin-related proteins, F-box and ubiquitination-associated genes, subtilase proteases, cytochrome P450s, glycosyltransferases, and transcriptional regulators. This functional bias suggests that gene family turnover is not random, but preferentially affects pathways involved in pathogen recognition, immune signaling, detoxification, and stress regulation. Such enrichment is consistent with repeated expansion of plant defense-related gene families (Roulin et al., 2013; Edger and Pires, 2009; Meyers et al., 2005) and with Red Queen dynamics, where co-evolutionary arms races with pathogens favor diversification of recognition and immune gene families (Van Valen, 1973). The enrichment of stress- and metabolism-related families further suggests that abiotic pressures—such as detoxification demands and environmental stress—also contribute to gene family turnover independently of biotic interactions Hanada et al. (2008).

The fate of duplicated copies further indicates that gene family expansion is a balance between retention, decay, and occasional innovation. Functional divergence was present but rarely dominated entire families, suggesting that neofunctionalization or subfunctionalization usually affects only subsets of retained duplicates (Force et al., 1999; Innan and Kondrashov, 2010). Pseudogenization was widespread, particularly in *A. lyrata*, consistent with relaxed constraint after duplication (Zhang, 2003; Scannell et al., 2007). Together, these patterns support a birth–death framework in which duplication generates functional redundancy, but long-term persistence depends on selection, dosage constraints, and genomic context (Kondrashov, 2012). Genome architecture appears to modulate these outcomes. The reduced chromosomal clustering and stronger TE association of duplicated-family genes in *A. lyrata* suggest that dispersed duplication, rearrangement, and TE-associated processes contributed more strongly to gene family evolution in this lineage. TEs can promote gene movement, alter regulatory contexts, and accelerate pseudogenization through insertion or ectopic recombination (Feschotte and Pritham, 2007; Bennetzen and Wang, 2014; Quesneville, 2020; Rebollo et al., 2012; Oliver and Greene, 2009). This provides a plausible explanation for the more variable duplication landscape in *A. lyrata*, whereas the lower TE burden and more constrained genome of *A. thaliana* may favor greater retention of intact duplicated copies (de la Chaux et al., 2012).

CNV within rapidly evolving families represents a recent layer of divergence superimposed on these deeper evolutionary dynamics. In *A. thaliana*, CNV shows a diffuse geographic and environmental structure, extending previous SNP- and CNV-based studies (Alonso-Blanco et al., 2016; Göktay et al., 2020; Kang et al., 2023; Zmienko et al., 2020; Lian et al., 2024). In *A. lyrata*, CNV is more tightly aligned with major phylogeographic divisions, consistent with known population structure (Scott et al., 2025; Hämälä et al., 2018b). Thus, CNV reflects both local ecological differentiation and demographic history, with the relative contribution of each differing between species.

Our results place previous CNV studies into a broader comparative framework. Earlier work often focused on selected loci, candidate families, or single-species analyses, showing that CNV contributes to adaptive, metabolic, and stress-related variation (Shirai et al., 2017; Wei et al., 2025; Wilson et al., 2025). Here, we show that these signals scale to genome-wide gene family landscapes and recur across related species, particularly in defense-related functional space. At the same time, the architecture of CNV–environment association differs: *A. thaliana* shows a broader, more polygenic climatic signal, whereas *A. lyrata* shows stronger coupling between CNV, phylogeographic structure, and genome turnover.

Overall, this study supports a model in which gene duplication provides raw material for environmental responsiveness, but the evolutionary outcome depends on functional context, TE-associated genome architecture, and lineage history. Defense- and stress-related families are repeatedly recruited into rapid turnover and CNV–environment associations, while duplicated copies are differentially retained, diversified, or lost across species. CNV therefore represents a flexible but context-dependent mechanism contributing to adaptation and genome evolution in *Arabidopsis*.

## 5 Conclusion

In summary, our study demonstrates that gene family evolution in *Arabidopsis* is governed by a combination of conserved birth–death dynamics and lineage-specific genomic processes. While most gene families show stable turnover, rapidly evolving families are consistently enriched for defense- and stress-related functions, indicating that CNV is preferentially concentrated in pathways mediating environmental responsiveness. By integrating long-read assemblies with a comparative, multi-species framework, we show that CNV patterns differ markedly between closely related species: *A. thaliana* exhibits a diffuse, polygenic CNV landscape with partially independent environmental associations, whereas *A. lyrata* shows stronger coupling between CNV, genome structure, and phylogeographic history. These differences are further shaped by genome architecture, particularly TE-associated processes that influence duplication, dispersal, and gene loss. Together, our results extend previous locus-specific studies by demonstrating that CNV–environment relationships emerge at the level of gene family networks and are modulated by lineage history and genomic context. This highlights CNV as a flexible but context-dependent mechanism contributing to genome evolution and ecological differentiation in plant species.

## Acknowledgment

This work was supported by the Deutsche Forschungsgemeinschaft (DFG, German Research Foundation) - Project ID 456082119 - TRR 341, in Cologne and Düsseldorf.

## Competing Interests

All authors declare no conflict of interest.

## Author Contributions

LR, TW, and PK conceived and designed the study. PK developed the analytical workflow, performed the analyses, and wrote the manuscript with input from all authors. YS carried out SNP calling and transposable element profiling. AG and LA generated and assembled the long-read sequencing data. LR and TW secured funding. All authors contributed to revisions and approved the final manuscript.

## Data Availability

The reference genomes used in the analysis are from *A. thaliana*: TAIR10: GCA_000001735.1 Cheng et al. (2017); *A. lyrata*: GCA_000004255.1 Hu et al. (2011); Rawat et al. (2015); *A. halleri*: [Arabidopsis_halleri v2.03, DOE-JGI](http://phytozome.jgi.doe.gov/); *A. alpina*: GCA_000733195.1; and *Carica papaya*: [Carica_papaya ASGPBv0.4](https://phytozome-next.jgi.doe.gov/info/Cpapaya_ASGPBv0_4) Ming et al. (2008); with their corresponding annotations. The long-read sequences of *A. thaliana* were downloaded from PRJNA777107, PRJNA779205 PRJNA834751, PRJNA1033522, PRJEB55353, PRJEB50694, and PRJEB55632 NCBI bio-projects with their corresponding meta-data. Genome assemblies of *A. thaliana* are also from above NCBI bioprojects. The *A. lyrata* long-read genome assemblies of *A. lyrata* are deposited in ENA under the project numbers PRJEB61074 and PRJEB96779, and will be released publicly upon acceptance (can be accessed on request before). The *A. lyrata* WGS resequencing data from Scott et al. (2025) used to generate SNPs can be accessed from ENA database under project numbers PRJEB67879 and PRJEB60410. All the scripts and intermediate data used in the analysis are archived in a GitHub repository and can be accessed at https://github.com/piyalkarum/GeneFamilyEvolution. All intermediate and processed data can be provided upon request.

## Supplementary Information (brief legends)

### Methods

- S1. Analysis of gene family expansion and contraction
- S2. Long-read sequencing and assembly preparation
- S3. Copy Number Variation Detection in Defense and Stress-Response related Gene Families using de novo Assemblies
- S4. SNP Dataset and Population Structure Assessment

### Figures

- Figure S1: Model selection for CAFE5 analysis of gene family evolution.
- Figure S2: Coefficient of variation (CV) in copy number variation across highly variable gene families in *Arabidopsis*.
- Figure S3A&B: Copy number variation in rapidly evolving gene families in *A. thaliana* and *A. lyrata*.
- Figure S4: PCA clustering of genetic divergence among populations of *A. thaliana* and *A. lyrata*.
- Figure S5: Combined PCA of *A. thaliana* and *A. lyrata* based on CNV structure of rapidly evolving gene families.
- Supplementary Figure S6: Functional category enrichment of gene families showing CNV– environment associations.

### Tables

- Table S0: Reference proteomes, genome assemblies, and their annotations used in this study.
- Table S1: Summary of gene co-localization in gene families that experienced significant rates of expansion and contraction.
- Table S2: Summary statistics of CNV assessment in all rapidly evolving gene families in *A*.*thaliana* (AT) and *A*.*lyrata* (AL).
- Table S3: *A. thaliana* long-read assemblies with meta data, assembly-wise gene family sizes, BUSCO summary, and TE summary.
- Table S4: *A. lyrata* long-read assemblies with meta data, assembly-wise gene family sizes, BUSCO summary, and TE summary.
- Table S5: Environmental variables used in the CNV-ENV analysis and corresponding RDA summary statistics.
- Table S6: GO enrichment assessment of rapidly-evolving gene families.
- Table S7&8(.1&.2): Gene structure and functional integrity assessment of all rapidly evolving gene families of *A. thaliana* and *A. lyrata*.
- Table S9: Assessment of technical and TE confounders on CNV variation.
- Table S10(.1&.2): SVD of covariance matrix representing population structure of the two species.
- Table S11: Functional description and categorization of all members of rapidly evolving gene families.

## Notes

### Competing Interest Statement

The authors have declared no competing interest.

### Summary of Updates

Briefly, the main changes are that we expanded the analysis to all rapidly evolving gene families instead of focusing only on defense- and stress-response-related genes. With this broader framework, defense- and stress-related gene families now emerge as significantly associated with environmental variables. In addition, we implemented a more thorough functional and evolutionary fate assessment, which replaces the traditional population genetic summary statistics used previously.

