## supplementary information for "Copy Number Variation: A Substrate for Plant Adaptation and Stress Response in Arabidopsis"

### S1. Analysis of gene family expansion and contraction

Gene families were inferred as orthogroups using OrthoFinder2 (Emms and Kelly, 2019), which defines an orthogroup as the complete set of genes descended from a single gene in the last common ancestor of the species considered. This definition aligns with the concept of a gene family as a set of homologous genes that have evolved from a common ancestor through speciation and/or duplication events. By including multiple closely related species and an outgroup, we ensured that orthogroup delineation reflects both recent and ancient gene duplications, improving the resolution of gene family boundaries.

We assessed the rate of gene family evolution in the selected Brassicaceae species using the curated reference proteomes of *A. thaliana*, *A. lyrata*, *A. halleri*, and *Arabis alpina* (see Tables S3&4 for details). We used *Carica papaya* as the outgroup, a member of the family Caricaceae (order Brassicales). The data was downloaded from Ensemble reference database (Martin et al., 2022). We only kept the primary proteins (longest transcript of a gene) for all the genes from these proteomes to avoid complications from variations of proteins in the gene family discovery.

In summary, OrthoFinder infers orthogroups using the original OrthoFinder algorithm (Emms and Kelly, 2015) where an all-vs-all BLAST is performed with a bit score to eliminate the bias caused by sequence length. The bit score is then used to group genes (i.e., orthogroups) based on a graph-based clustering approach called MCL (Markov Clustering Algorithm). An unrooted gene tree is inferred for each orthogroup using DendroBLAST. The unrooted species tree is inferred from this set of unrooted orthogroup trees using the STAG algorithm. This STAG species tree is then rooted using the STRIDE algorithm by identifying high-confidence gene duplication events in the complete set of unrooted orthogroup trees. The rooted species tree is used to root the orthogroup trees. Orthologs and gene duplication events are inferred from the rooted orthogroup trees by a novel hybrid algorithm that combines the “species-overlap” method and the duplication-loss-coalescent model. Finally, comparative statistics are calculated.

All orthogroups with high confidence and resolved trees from OrthoFinder were used as gene families for assessing the evolutionary rates of gene families among species. The evolutionary rates based on the gene Birth and Death (BD) model of gene family evolution is calculated using the CAFE 5 software (Hahn et al., 2005; Mendes et al., 2020). The algorithm in CAFE uses the number of genes in each species for each gene family and a time-calibrated species tree to determine the gene family expansion and contraction rate $\lambda$ parameter, which represents the rate of gene gain (birth) and loss (death) per gene per million years. CAFE 5 uses a likelihood ratio test (LRT) to find the best model and Monte Carlo simulations to calculate expected family sizes using the estimated $\lambda$ parameter. A chi-square test determines the significance.

$$\Delta[L] = 2(logL_1 - logL_0)$$

$L_0$ - likelihood of the null model ($\lambda = 0$)
$L_1$ - likelihood of the alternative model ($\lambda > 0$)
$\Delta[L]$ - likelihood ratio test statistic

In summary, CAFE 5 analyzes changes in gene family size in a way that accounts for phylogenetic history and provides a statistical foundation for evolutionary inferences. The program uses the birth and death process to model gene gain and loss across a user-specified phylogenetic tree. This enables the estimation of an evolutionary rate for gene family evolution as a global (single $\lambda$ values) or local (multiple $\lambda$ values) rate. The latter allows for adding more gamma distribution parameters to find the best fit model considering varying levels of evolutionary rates in different gene families.

In order to find the best $\lambda$ that describes the evolutionary rates among the studied species, we ran CAFE 5 with gamma parameters varying from $k = 1$ (base model) to $k = 5$ (maximum allowed for five species). The model outputs are log-likelihood and $\lambda$ estimations for each model. We used both the log-likelihood and AIC/BIC to determine the best model without overfitting the data.

An error model was run to account for genome annotation artifacts and the resulting error factor was used to rerun the base CAFE models. The error factor was 0.023207 and it did not significantly change the $\lambda$ values of the base model, indicating that the genome annotations are considerably accurate.

We initially used a maximum p-value of 0.05 to find the significantly fast evolving gene families under the base model, which was later refined using a more stringent p-value of 0.01. The gene functions of the members of these gene families were predicted as explained below.

### S2. Long-Read Sequencing and *de novo* Assembly Preparation of *A. lyrata*

High-molecular-weight genomic DNA was extracted from leaf tissues of 24 *A. lyrata* individuals representing the species' distribution range. The extracted DNA was used for long-read sequencing on the PacBio Sequel II platform. The raw PacBio FASTQ reads were assembled with Mabs (Schelkunov, 2023) and Hifiasm (Cheng et al., 2021), using the *Brassicales_odb*10 lineage dataset from BUSCO (Manni et al., 2021; Kriventseva et al., 2019) to guide the assembly optimization. The best assembly produced by Mabs was scaffolded using the `scaffolds` function of RagTag (Alonge et al., 2019), with the *A. lyrata* NT1 reference genome as the guide (https://figshare.com/articles/dataset/NT1_genome_assembly/22285429?file=39633610). The resulting RagTag scaffolding were further manually curated for accuracy. Long inversions relative to the NT1 scaffolds were identified and visualized using synteny plots generated by GENESPACE (version 1.4) (Lovell et al., 2022). Manual curation and correction of large-scale misassemblies and orientation errors were performed based on these synteny visualizations. Sequencing information and the quality statistics are in Tab. S4.

### S3. Detailed CNV detection pipeline for gene families

Copy number variation (CNV) across rapidly evolving gene families was inferred using an assembly-based, BLAST-driven pipeline designed for long-read *de novo* genome assemblies. The workflow integrates reference gene extraction, sequence similarity searches, clustering of genomic hits, and post-processing to resolve gene copies at the locus level.

**Reference gene set construction:** Gene families (orthogroups) were defined based on OrthoFinder2 clustering. For each orthogroup, corresponding *A. thaliana* reference gene models (Araport11 annotation) were extracted. Gene coordinates were obtained from the GFF3 annotation, and genomic sequences were retrieved from the TAIR10 reference genome using `Biostrings`. For each gene family, all member gene sequences were written to FASTA files, forming the query reference set. To ensure comparability across assemblies, only nuclear gene families were retained; orthogroups associated with non-chromosomal elements (e.g., mitochondrial genes) were excluded. Gene lengths were calculated for all reference sequences and used in downstream filtering steps.

**Sequence similarity search:** Reference gene sequences were queried against each long-read assembly using `blastn` (BLAST+ v2.9; Camacho et al., 2009) in pairwise mode with custom parameters (initial: `word-size=9, e-value=`$1e^{-5}$ for genes $>$ 500bp, $1e^{-2}$ for gene $<$ 500bp; confirmation pass: `word-size=11, e-value=`$1e^{-20}$). Searches were performed for each gene family against each assembly, and results were retained in tabular format including alignment coordinates, percent identity, and alignment length. This approach allows recovery of both full-length and partial matches, accommodating structural variation and assembly fragmentation.

**Filtering and clustering of BLAST hits:** Raw BLAST hits were filtered to retain only alignments with $\geq$90% sequence identity. For each gene and assembly, hits were grouped by scaffold and clustered based on genomic coordinates using a custom clustering procedure (`gSoup::find_blast_clusters`). Clusters were defined such that the genomic span of merged hits did not exceed 1.3$\times$ the reference gene length, thereby preventing artificial merging of adjacent loci. For each cluster, genomic intervals were reconstructed by taking the minimum and maximum coordinates of grouped hits. Candidate loci were retained only if their length exceeded 50% of the corresponding reference gene length, thereby excluding short fragments and spurious matches, including those arising from transposable elements or low-complexity regions. Each retained cluster was treated as a putative gene copy, and copies were enumerated sequentially within each assembly and gene.

**Resolution of overlapping loci:** To avoid overestimation of copy number due to redundant or ambiguous assignments, overlapping loci across different reference genes within the same orthogroup were further filtered. Genomic intervals overlapping by more than 70% were considered redundant ("pseudo-duplicates"). In such cases, only a single representative locus was retained, effectively collapsing overlapping signals arising from closely related paralogs or sequence similarity among gene family members.

**Estimation of copy number and CNV metrics:** For each gene family and assembly, copy number was quantified by counting the number of non-overlapping loci per gene. Gene-level counts were then aggregated to the family level. For each assembly, the following metrics were computed: **(1) Gene count:** number of detected genes with at least one copy. **(2) Gain:** number of genes with copy number $>$ 1, **(3) Loss:** number of genes with copy number $<$ 1 (i.e., absent relative to the reference set), **(4) Gain–loss count:** number of genes deviating from single-copy expectation, and **(5) Total gene count:** sum of all detected copies within the gene family. These metrics were calculated for all assemblies and gene families, resulting in a matrix of gene family copy number profiles across individuals.

**Construction of CNV matrices:** To facilitate downstream analyses, gene family copy number matrices were constructed by reshaping the data into wide format, with gene families as rows and assemblies as columns. Missing values were replaced with zeros, reflecting absence of detected copies. These matrices formed the basis for within-species CNV analyses, including variance estimation, multivariate ordination, and environmental association testing.

**Rationale and limitations:** This pipeline leverages the high contiguity and structural accuracy of long-read assemblies to directly resolve gene copy architecture at the sequence level. The BLAST-based approach enables detection of both intact and partial gene copies without reliance on predefined gene annotations in target assemblies. However, the method assumes that sequence similarity remains sufficiently high (≥90%) to detect homologous loci, which may limit detection of highly diverged copies. In addition, clustering thresholds (1.3× gene length; 50% coverage; 70% overlap) represent empirically chosen parameters that balance sensitivity and specificity but may influence copy number estimates in complex genomic regions. Despite these limitations, the approach provides a robust and transparent framework for gene family–focused CNV inference across large numbers of assemblies.

### S4. SNP Dataset and Population Structure Assessment

To determine the population structure among populations included in the CNV analysis, we conducted the following steps:

1) For *A. thaliana*, we downloaded long-read whole-genome sequencing data for 128 individuals representing the populations analyzed (Table S3) from the NCBI SRA database. The dataset comprised PacBio Sequel II HiFi reads and Oxford Nanopore Technologies (ONT) reads. All reads were mapped to the *A. thaliana* TAIR10 reference genome using `minimap2` (version 2.26; Li, 2018) with presets optimized for PacBio and ONT data (`-x map-hifi` and `-x map-ont`, respectively). Variant calling was performed with `bcftools mpileup` and `bcftools call` (version 1.18; Li, 2011), which efficiently handle both long-read platforms. The resulting per-sample VCF files were merged into a single, concatenated VCF, which was annotated with `SnpEff` (version 5.1; Cingolani et al., 2012) using the TAIR10 `.gff3` annotation file.

From the annotated variants, we extracted putatively neutral sites, defined here as SNPs located in intronic, intergenic, and four-fold degenerate coding positions, yielding 677,120 variable sites. To minimize linkage disequilibrium effects, we generated 10 independent subsets of 10,000 randomly selected SNPs and used them to compute an average covariance matrix among individuals.

2) For *A. lyrata*, we used the whole-genome SNP dataset from Scott et al. (2025), which comprises 1,018 Illumina short-read sequences representing 58 populations. From this dataset, we selected at least five individuals per population corresponding to those used in the CNV analysis. Putatively neutral sites were extracted from the annotated VCF files following the same criteria as for *A. thaliana*, resulting in 657,524 variable sites. As above, we generated 10 random subsets of 10,000 SNPs each to compute an average covariance matrix.

The covariance matrix assumes that neutral genetic distances among population pairs reflect evolutionary history and shared ancestry (Gautier, 2015). Population structure for each population was summarized by applying singular value decomposition (SVD) to the covariance matrix, and the resulting per-population values were used to correct for population structure in RDA and family-wise univariate analysis of CNV–environment associations (see Materials and Methods). Heatmaps and PCAs of the covariance matrices are shown in Fig. S3 (also see Table S10 for SVD of genetic diversity values).

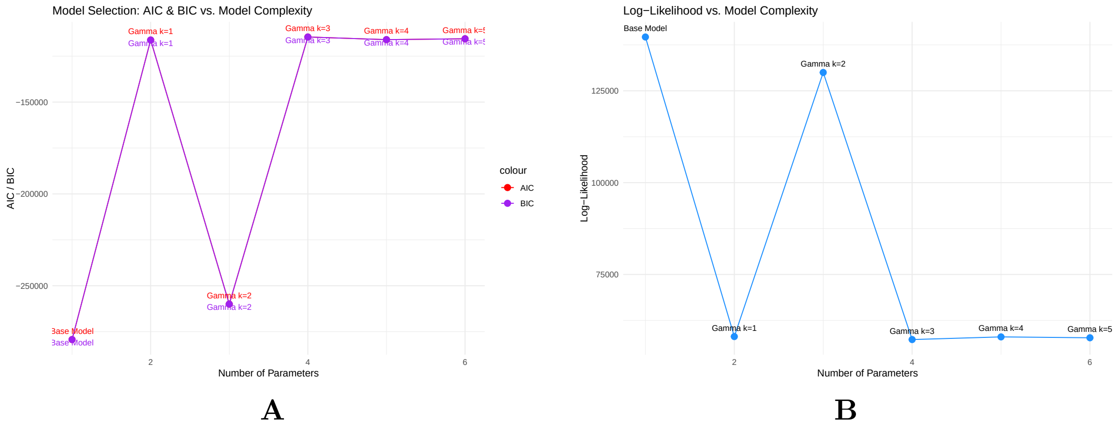


Figure S1: **Model selection for CAFE5 analysis of gene family evolution.** Model performance was evaluated across a range of models using different values of the gamma ($\gamma$) parameter, which allows the birth-death rate ($\lambda$) to vary across gene families. In CAFE5, $\gamma$ controls the number of discrete rate categories used to capture rate heterogeneity: higher $\gamma$ values allow more flexibility by modeling gene families as evolving under different $\lambda$ values, thus accommodating lineage- or function-specific evolutionary dynamics. Panel A shows Akaike Information Criterion (AIC) and Bayesian Information Criterion (BIC) values for models with increasing $\gamma$ values. Panel B shows the corresponding log-likelihood scores. Although models with larger $\gamma$ values typically achieve higher log-likelihoods by better fitting the observed data, this comes at the cost of increased model complexity. Both AIC and BIC penalize complexity, and in our case, they support the base model ($\gamma = 1$) as the best-fitting model, balancing goodness-of-fit with parsimony. This indicates that gene family turnover across the dataset is adequately captured by a single-rate model without requiring more complex multi-rate models.

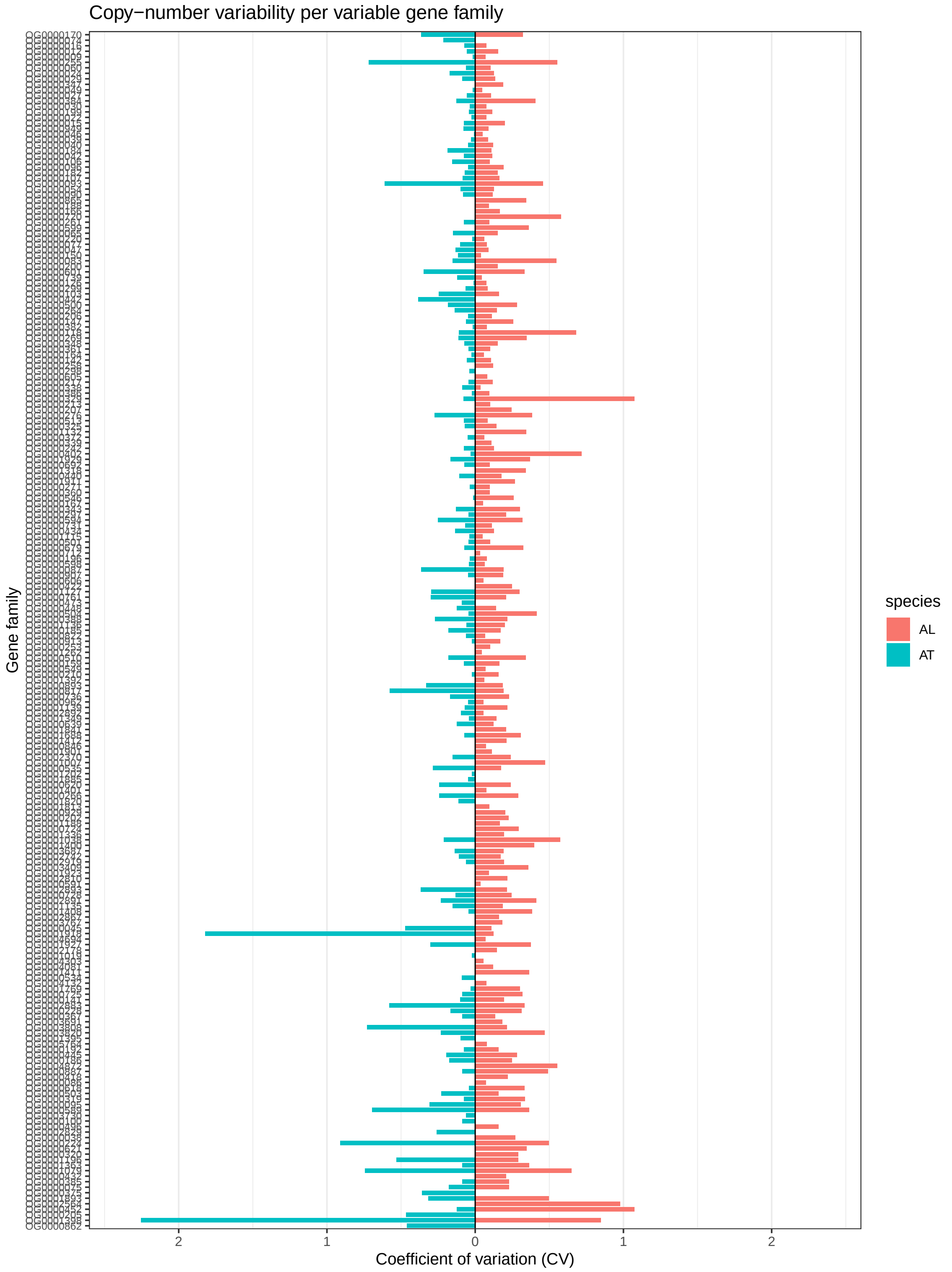


Figure S2: **Coefficient of variation (CV) in copy number variation across highly variable gene families in *Arabidopsis*.** The bars on left and right show the distribution of gene copy numbers across assemblies for *A. thaliana* (AT; left) and *A. lyrata* (AL; right). X-axis is the CV and the Y-axis is the orthogroup ID (only families variable in both species are plotted; i.e., CNV present families in both species).

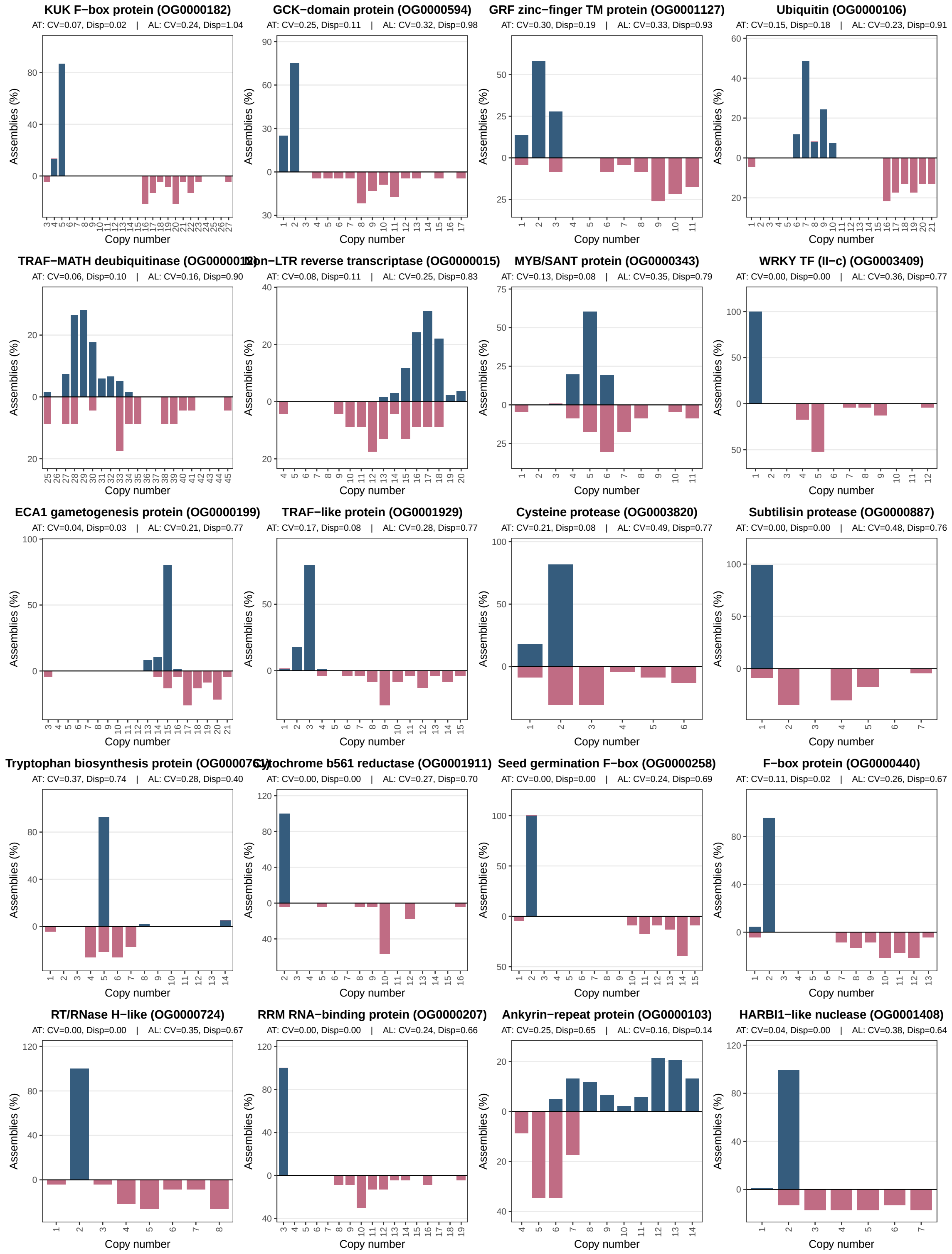


Figure S3A: **Distribution of copy number variation across highly variable gene families in *Arabidopsis.*** Each panel represents one gene family (top 30 ranked by dispersion across species), showing the distribution of gene copy numbers across assemblies for *A. thaliana* (AT; upward bars) and *A. lyrata* (AL; downward bars). Bar heights correspond to the percentage of assemblies exhibiting a given copy number, with species-specific scaling applied within each family. Panel titles indicate the short functional annotation of each gene family followed by the orthogroup ID. Subtitles report the coefficient of variation (CV) and dispersion (variance-to-mean ratio) for each species. These plots highlight pronounced heterogeneity in CNV distributions among gene families and reveal contrasting patterns of copy number variability between the two species.

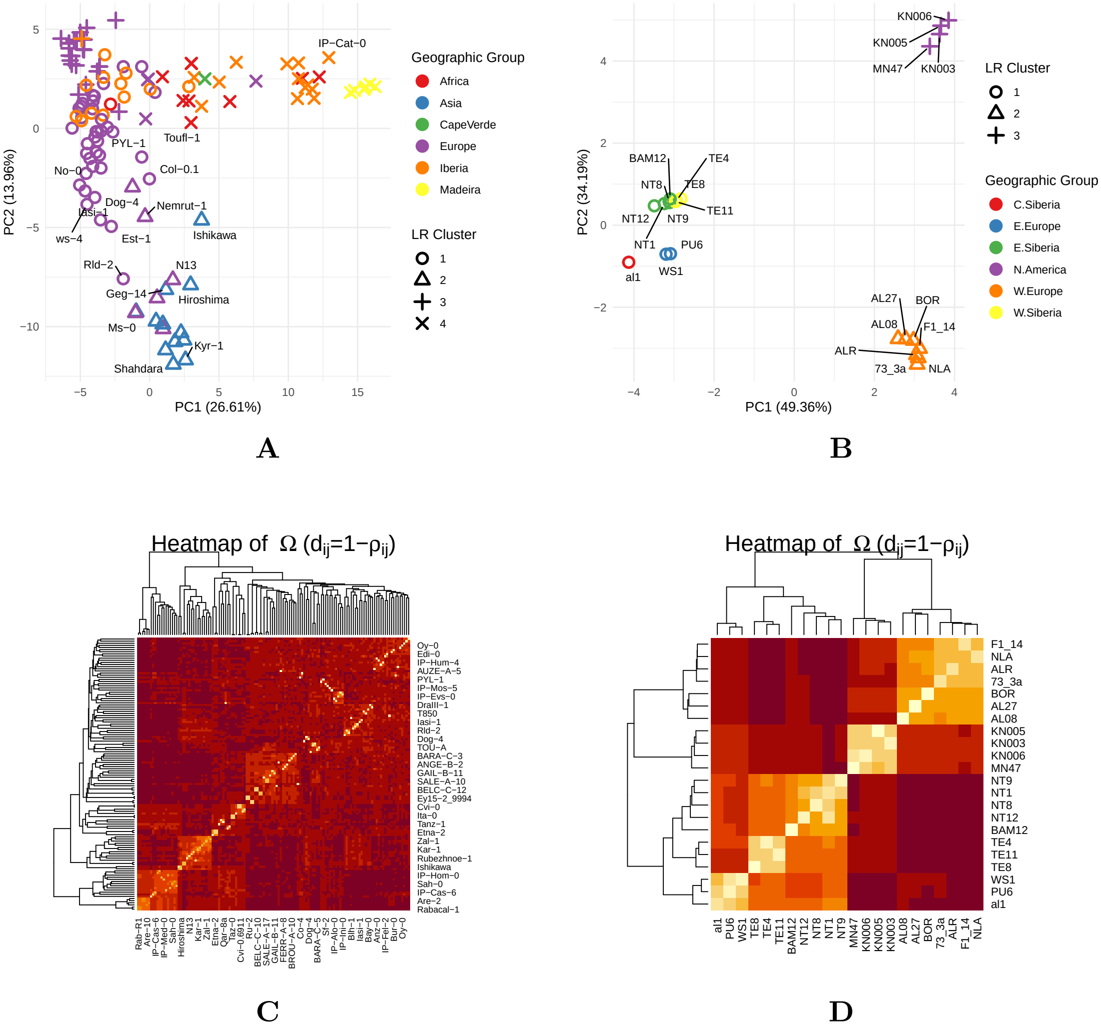


Figure S4: **PCA clustering of genetic divergence among populations of *A. thaliana* (A) and *A. lyrata* (B).** The PCA was performed on the covariance (Ω) matrix derived from putatively neutral loci (four-fold degenerate sites, introns, and intergenic regions), following (Gautier, 2015), where genetic distance between two populations is assumed to co-vary based on their shared history. PCA axes are the single value decomposition (SVD) of pairwise covariance. The plot characters indicate the genetic clusters of populations, with colors representing different geographical regions. A 3D plot of *A. thaliana* with first three PC axes is included in the supplementary plot '3D_SVD_plot.html'. Genetic clusters were determined using hierarchical clustering of the covariance matrix. **C&D** show the SVD heatmaps of Ω matrices for *A. thaliana* and *A. lyrata*, respectively, where color intensity indicates pairwise population similarity: darker reds denote lower covariance (greater divergence), whereas yellow to light-yellow (flame-like colors) indicate higher covariance (greater similarity).

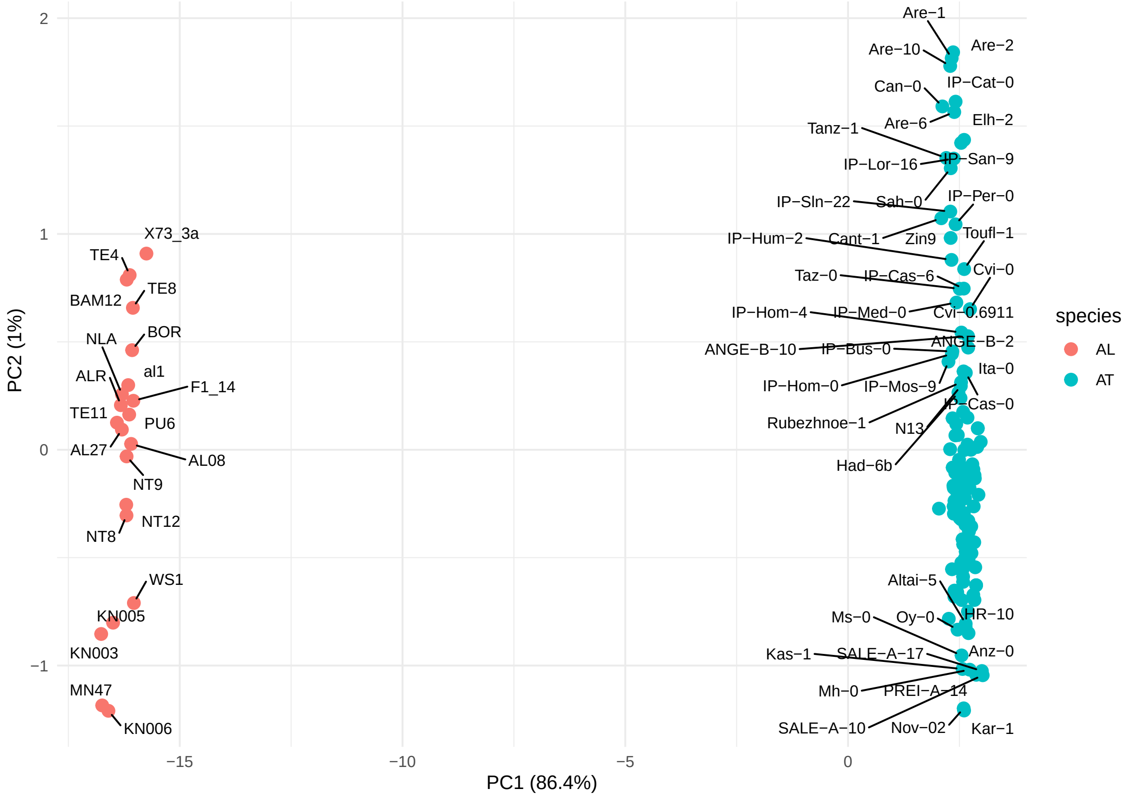


Figure S5: **Combined PCA of *A. thaliana* and *A. lyrata* based on CNV structure of rapidly evolving gene families.**

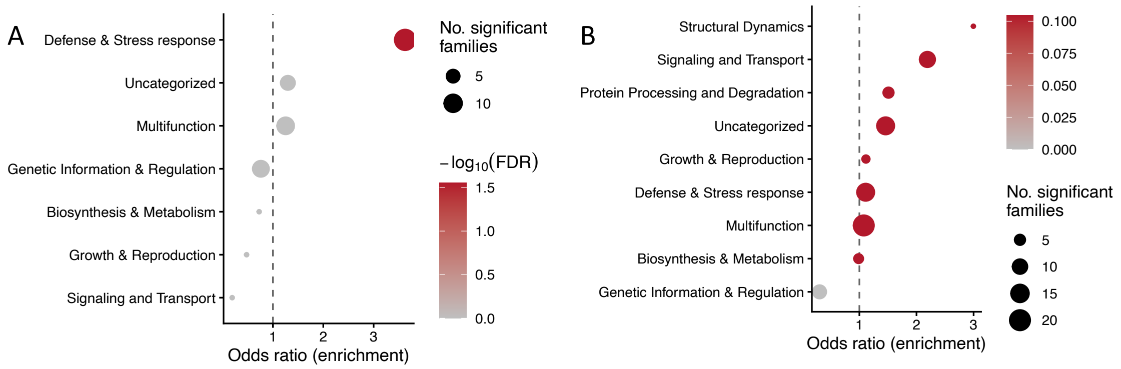


Figure S6: **Functional category enrichment of gene families showing CNV–environment associations.** Functional enrichment results from family-wise univariate analyses are shown for *A. thaliana* (A) and *A. lyrata* (B). Points represent functional categories, with enrichment expressed as odds ratios, point size indicating the number of significant gene families, and color corresponding to statistical significance (-$\log_{10}$ FDR). Defense- and stress-related as well as multifunctional categories show the strongest enrichment among environmentally associated gene families.

Table S0: Reference proteomes, genome assemblies, and their annotations used in this study.

| Species | Ecotype | Source | Assembly | Annotation | Genes | N50 (Mb) | BUSCO (%) |
|---|---|---|---|---|---|---|---|
| *A. thaliana* | Col-0 | Phytozome | TAIR9 | Araport11 | 27,655 | 23.5 | 99.3 |
| *A. lyrata* | MN47 | Phytozome | v1 | v2.1 | 31,073 | 24.5 | 98.2 |
| *A. halleri* | Lan3.1 | Phytozome | v2.03 | v2.1.0 | 28,722 | 23.0 | 94.0 |
| *A. alpina* | Pajares | arabis-alpina.org | V4 | V4 | 30,729 | 0.79[a] | 79.8 |
| *C. papaya* | — | Phytozome | r.Dec2008 | ASGPBv0.4 | 27,769 | 1.18 | 72.0 |

*Genes* = number of protein-coding genes; *BUSCO* = Embryophyta lineage (OrthoDB v9).
[a] Scaffold-level assembly (11,244 scaffolds; 309 Mb total assembled).

Table S1: **Summary of gene localization in gene families that experienced significant rates of expansion and contraction**. *clustered* indicates the number of gene families with members on the same chromosome, while *dispersed* indicates the number of gene families with members on different chromosomes.

**Table S1A.** Gene family localization summary

| Localization | *A. alpina* | *A. halleri* | *A. thaliana* | *A. lyrata* |
|---|---|---|---|---|
| **Clustered** | 46 | 42 | 46 | 28 |
| **Dispersed** | 124 | 113 | 108 | 170 |

**Table S1B.** Fisher's Exact Test $p$-values

| | *A. alpina* | *A. halleri* | *A. thaliana* |
|---|---|---|---|
| ***A. halleri*** | 1.0000 | – | – |
| ***A. thaliana*** | 0.7468 | 0.7468 | – |
| ***A. lyrata*** | 0.0060 | 0.0060 | 0.0022 |
